# A single origin of animal excretory organs

**DOI:** 10.1101/2020.11.15.378034

**Authors:** Ludwik Gąsiorowski, Carmen Andrikou, Ralf Janssen, Paul Bump, Graham E. Budd, Christopher J. Lowe, Andreas Hejnol

## Abstract

Excretion is an essential physiological process, carried out by all living organisms regardless of their size or complexity(1–3). Most animals, which include both protostomes (e.g. flies, flatworms) and deuterostomes (e.g. humans, sea urchins) (together Nephrozoa(4, 5)), possess specialized excretory organs. Those organs exhibit an astonishing diversity, ranging from units composed of just three distinct cells (e.g. protonephridia) to complex structures, built by millions of cells of multiple types with divergent morphology and function (e.g. vertebrate kidneys)(6, 7). Although some molecular similarities between the development of kidneys of vertebrates and the regeneration of the protonephridia of flatworms have been reported(8, 9), the molecular development of nephrozoan excretory organs has never been systematically studied in a comparative context(6). Here we show that a set of highly conserved transcription factors and structural proteins is expressed during the development of excretory organs of six species that represent major protostome lineages and non-vertebrate deuterostomes. We demonstrate that the molecular similarity witnessed in the vertebrate kidney and flatworm protonephridia(8) is also seen in the developing excretory organs of other Nephrozoa. In addition, orthologous structural proteins forming the ultrafiltration apparatus are expressed in all these organs in the filter-forming cells. Our results strongly suggest that excretory organs are homologous and are patterned by the conserved set of developmental genes. We propose that the last common nephrozoan ancestor possessed an ultrafiltration-based, ciliated excretory organ, a structure that later gave rise to the vast diversity of extant excretory organs, including the human kidney.

**Significance statement:** Most of the bilaterally symmetrical animals excrete through specialized excretory organs, such as kidneys and nephridia. However, due to the morphological diversity of these organs, it remains unknown whether those structures evolved from a common ancestral organ or appeared several times independently during evolution. In order to answer the question about the origin of excretory organs we investigated the molecular pathways and structural genes involved in the development of nephridia in 6 animal species representing major evolutionary lineages. We show that diverse excretory organs share an ancient molecular patterning and structural molecules. Our results provide strong evidence that all excretory organs originated from a single, simple organ that performed urine production by ultrafiltration in deep geological past.

## Main text

Excretion, the removal of metabolic waste products, is indispensable for any living organism. Most organisms excrete through the entire body surface(1, 2), however specialized excretory organs emerged during animal evolution (Fig. 1a). These organs are thought to be one of the key evolutionary innovations of the emergence of complex body plans(4, 5), and are believed to facilitate the conquest of new habitats, such as freshwater and terrestrial environments(3). Animals with specialized excretory organs form a group called Nephrozoa, which includes both protostomes (e.g. flies, flatworms) and deuterostomes (e.g. humans, sea urchins) (Fig. 1a), comprising most of animal biodiversity(4, 5, 10). The diverse nephrozoan excretory organs can be grouped into secretory organs (e.g. Malpighian tubules of insects), in which primary urine is produced by the means of active, transcellular transport(3, 6), and organs that are based on the principle of ultrafiltration (UF). The latter is a pressure-driven physiological process in which the body fluid (e.g. blood) of the organism is filtered through an extracellular filter to produce primary urine(3, 6, 11-14). UF-based organs include human kidneys, but also protonephridia and metanephridia, present in numerous invertebrates(3, 6, 11–14). These extracellular filters share ultrastructural and molecular properties, even between distantly related animals with divergent morphologies of their excretory organs(9), which might suggest a common evolutionary origin of UF(3, 11–14). Two main hypotheses on the evolution of the excretory organs have been proposed based on their comparative morphology(3). In the first(13, 14), all the UF-based excretory organs are homologous and differences in their architecture depend solely on the animal body size (protonephridia in small animals and metanephridia in larger ones). The second hypothesis proposes that protonephridia represent the ancestral organs that were replaced by the metanephridia and kidneys, several times independently, as a consequence of the development of the secondary body cavities (i.e. the coelom)(11).

**Figure 1.**
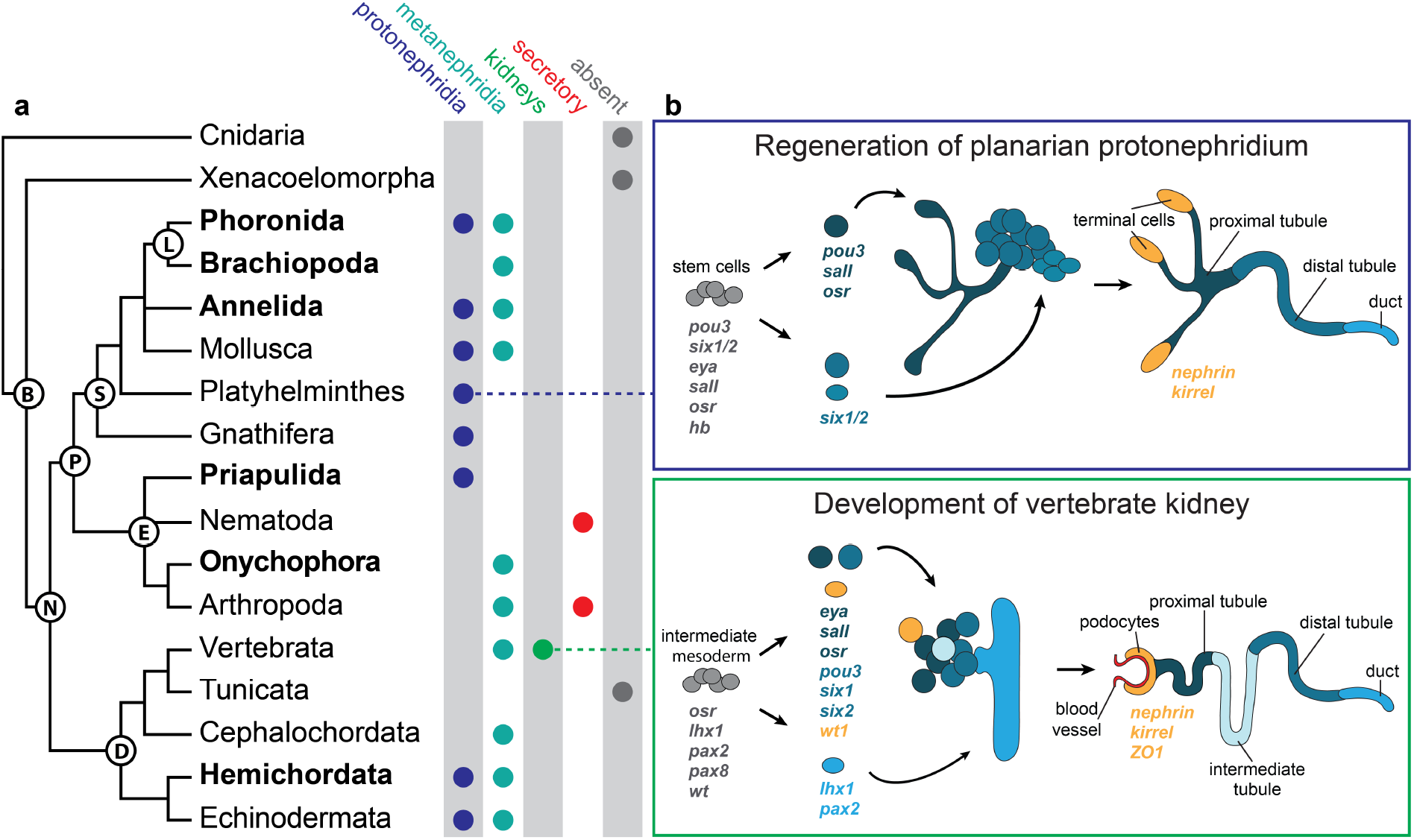
Phylogenetic distribution of excretory organs and their molecular patterning. **a**, Most of the Nephrozoan clades possess excretory organs based on the principle of ultrafiltration (UF) (protonephridia, metanephridia and kidneys). Specialized secretory-excretory organs are found in nematodes and some arthropods. Clades investigated in this study are marked in bold. **b**, Homologous transcription factors involved in regeneration of planarian protonephridium and development of the vertebrate kidney, in both organs homologous structural genes form the UF site (terminal cells and podocytes, respectively). Abbreviations: B, Bilateria; D, Deuterostomia; E, Ecdysozoa; L, Lophophorata; N, Nephrozoa; P, Protostomia; S, Spiralia.

Despite the importance of excretory organs for animal evolution, relatively little is known about the molecular basis of their development: the developmental genetic interactions of UF-based excretory organs have been so far described only for the vertebrate kidney (Fig. 1b)(6). Interestingly, some of the transcription factors (TFs) (*eya, six1/2, pou3, sall, osr*) involved in the development of the vertebrate kidney are also expressed during the regeneration of the flatworm protonephridia (Fig. 1b)(8). These similarities might suggest homology between kidneys and protonephridia(8). However, even though sets of conserved regulatory molecules might imply putative common ancestry of a structure between species(15, 16), such inference becomes problematic when the homology is based on findings in only two distantly related species(17). Evolutionary comparisons require broader sampling of multiple intermediate evolutionary lineages that could bridge the evolutionary distance(17, 18). Therefore, a comparison of gene expression during the development of multiple, divergent excretory organs of a number of Nephrozoa is desired to test their homology. If this homology is found, it will ultimately allow the reconstruction of the ancestral nephrozoan excretory organs, which potentially gave rise to the vastly diverse structures such as protonephridia, metanephridia and the human kidney.

## Results and discussion

### Molecular development of spiralian nephridia

In order to study gene expression during nephridial development in two closely related species, with divergent morphology of their excretory organs, we first investigated two animals – the phoronid *Phoronopsis harmeri* and the brachiopod *Terebratalia transversa.* Phoronids and brachiopods (together lophophorates) belong to the large animal group called Spiralia (Fig. 1a), which also includes flatworms, molluscs and annelids (segmented worms)(4, 10). We studied the expression of *eya, six1/2, pou3, sall, hb* and *osr,* genes known to be involved in the regeneration of planarian protonephridia(8), as well as *lhx1/5,* a homologue of the vertebrate gene *lim1*, indispensable for the formation of nephric tubules in vertebrates(19, 20).

Phoronids develop through a long-lived larva, which possess a pair of ciliated protonephridia (Figs. 2a, S1a)(21, 22). During metamorphosis this type of excretory organ is only partially lost and gets remodelled and incorporated into another type of excretory organ, the definitive adult metanephridium(21, 22). We detected expression of all of our candidate genes in the developing protonephridia (Figs. 2b, S1a), with the exception of *osr,* which instead was exclusively expressed in the sphincters of the digestive tract (Fig. 3a). Double *in situ* hybridization showed that most of the TFs are co-expressed in the identical cells of the protonephridial rudiments at the pre-tentacular larval stage (Fig. S1b), similar to what was demonstrated for the progenitor cells, from which protonephridia regenerate in flatworms(8).

**Figure 2.**
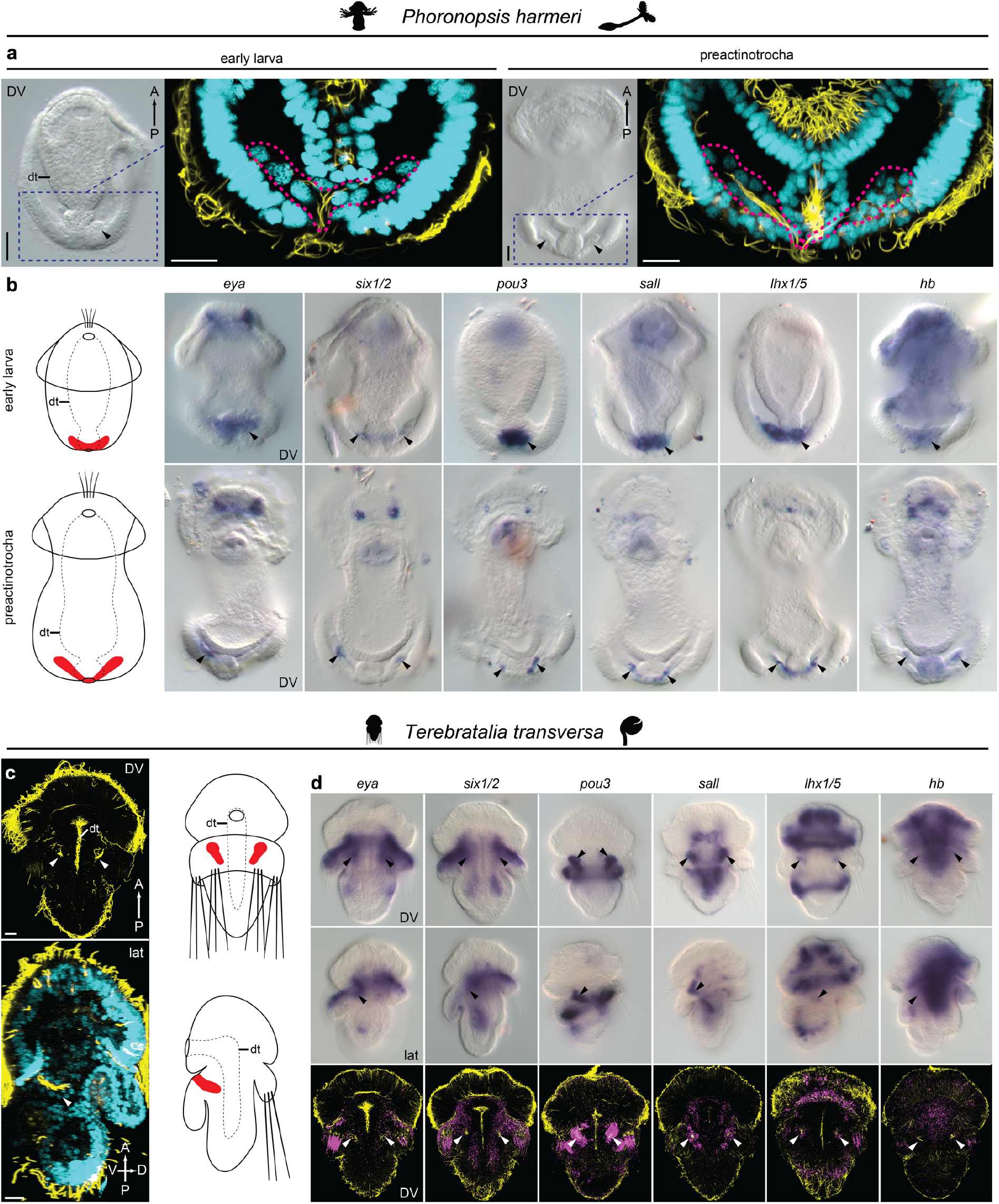
Expression of the nephridia-related transcription factors in Lophophorates. **a**, A pair of protonephridia (arrowheads) develops in the posterior end of the phoronid larva, ciliated cells forming in each protonephridium are outlined with magenta dotted lines. **b,** Transcription factors expressed in the developing protonephridia of *P. harmeri* (arrowheads), **c**, A pair of ciliated nephridial rudiments (arrowheads) is present in the middle portion of the larval *T. transversa.* **d**, Transcription factors are expressed in various structures, including the area where nephridia develop (arrowheads). Abbreviations: A, anterior; D, dorsal; dt, digestive tract; vv, ventral view; lat, lateral view; P, posterior; V, ventral. DAPI stained cell nuclei are in cyan and acetylated tubulin immunoreactivity is in yellow. Nephridia on the schematic drawings are marked in red. Scale bars, 20 μm.

**Figure 3.**
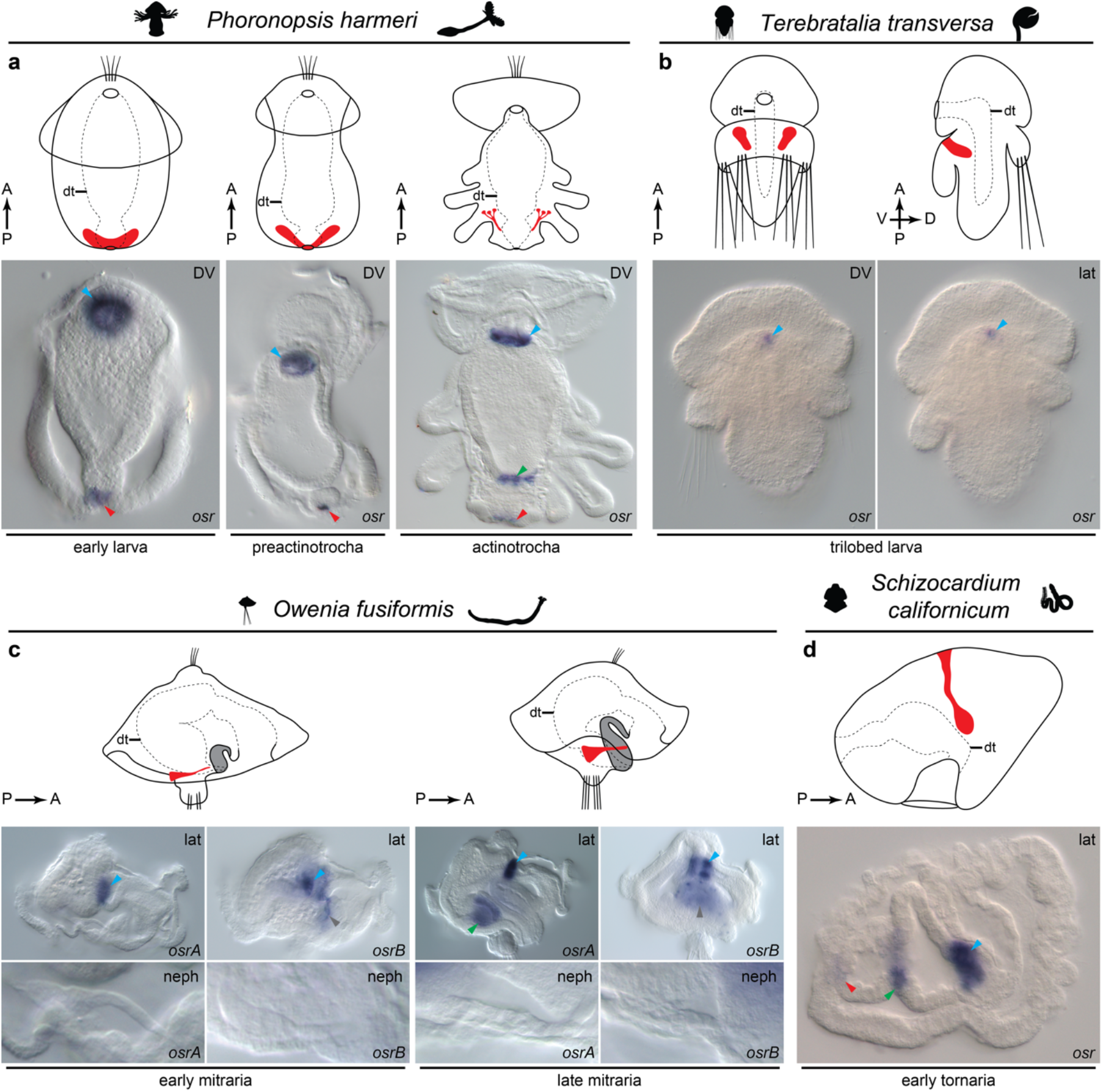
Expression of *osr* in larval Spiralia and hemichordate. **a**, In the earlier phoronid larvae *osr* is expressed in the esophageal sphincter (blue arrowheads) and around anus (red arrowheads); in the later actinotrocha the third expression domain appears at the midgut-hindgut transition (green arrowhead). **b**, *Osr* is expressed in the anterior digestive tract (blue arrowhead) of the larval *T. transversa.* **c**, *O. fusiformis* possesses two paralogues of *osr*: *osrA* is expressed in the esophageal sphincter (blue arrowhead) in early mitraria and in the esophageal sphincter (blue arrowhead) and posterior stomach (green arrowhead) in late mitraria, while *osrB* is expressed in both larval stages in the esophageal sphincter (blue arrowheads) and in the worm rudiment (grey arrowheads). **d**, *Osr* has three expression domains in the early larva of *S. californicum*: in the stomodaeum-midgut transition (blue arrowhead), midgut-hindgut transition (green arrowhead) and in the developing anal region (red arrowhead). Abbreviations: A, anterior; D, dorsal; dt, digestive tract; vv, ventral view; lat, lateral view; neph, magnified nephridium; P, posterior; V, ventral. Nephridia on the schematic drawings are marked in red.

In contrast to phoronids, brachiopods develop through a rather short-lived larval stage(23). Those larvae possess a pair of simple ciliated structures interpreted as larval nephridial rudiments (Fig. 2c)(23), which later give rise to the adult metanephridia shortly after metamorphosis (Fig. S2a)(24). We detected expression of *eya, six1/2, pou3, sall, lhx1/5* and *hb* in those larval nephridial rudiments (Fig. 2d), which persisted (with the exception of *hb*) in the juvenile metanephridia after metamorphosis (Fig. S2b). As in phoronids, we did not detect expression of *osr* in the nephridial tissue; instead, the gene was expressed in the anterior digestive tract in both larvae and juveniles (Figs. 3b, S2b). These data suggest that the same developmental regulatory program used for patterning of protonephridia (in both the planarians(8) and phoronids), seems to be deployed for the brachiopod metanephridia, despite their different morphology.

While most spiralians excrete either by simple protonephridia or typical metanephridia (Fig. 1a)(3, 11), some of them evolved more divergent types of excretory organs. To test whether these more aberrant nephridial types also express the same genes, we decided to study the annelid *Owenia fusiformis.* As a larva, *O. fusiformis* possesses distinct excretory organs (Fig. 4a) that differ greatly from the excretory organs of other spiralians, and based on their morphology and ultrastructure they were described as “deuterostome-like nephridia”(25). In contrast to the previous reports(25, 26), we found that larval protonephridia contribute to the adult excretory organs (Fig. S3), while structures interpreted previously as metanephridial rudiments(27) are non-ciliated (Fig. S3) and probably represent prospective tube-secreting glands(28). *Eya, six1/2, sall* and *hb* were all expressed in the protonephridia of both early and advanced larvae (Fig. 4b), while *pou3* expression was detected in the nephridial duct only at the late larval stage (Fig. 4b). Out of the four paralogues of *lhx1/5* present in the transcriptome of *O. fusiformis,* one was detected in the developing nephridia (Fig. 4b), while neither of the two *osr* paralogues was expressed in the nephridial tissues (Fig. 3c) (as in the phoronid and brachiopod). Instead, they were expressed in the sphincters of digestive tract and the epidermis of the adult worm rudiment (Fig. 3c). Altogether these results show that even peculiar, lineagespecific spiralian excretory organs deploy the same conserved set of molecular developmental TFs.

**Figure 4.**
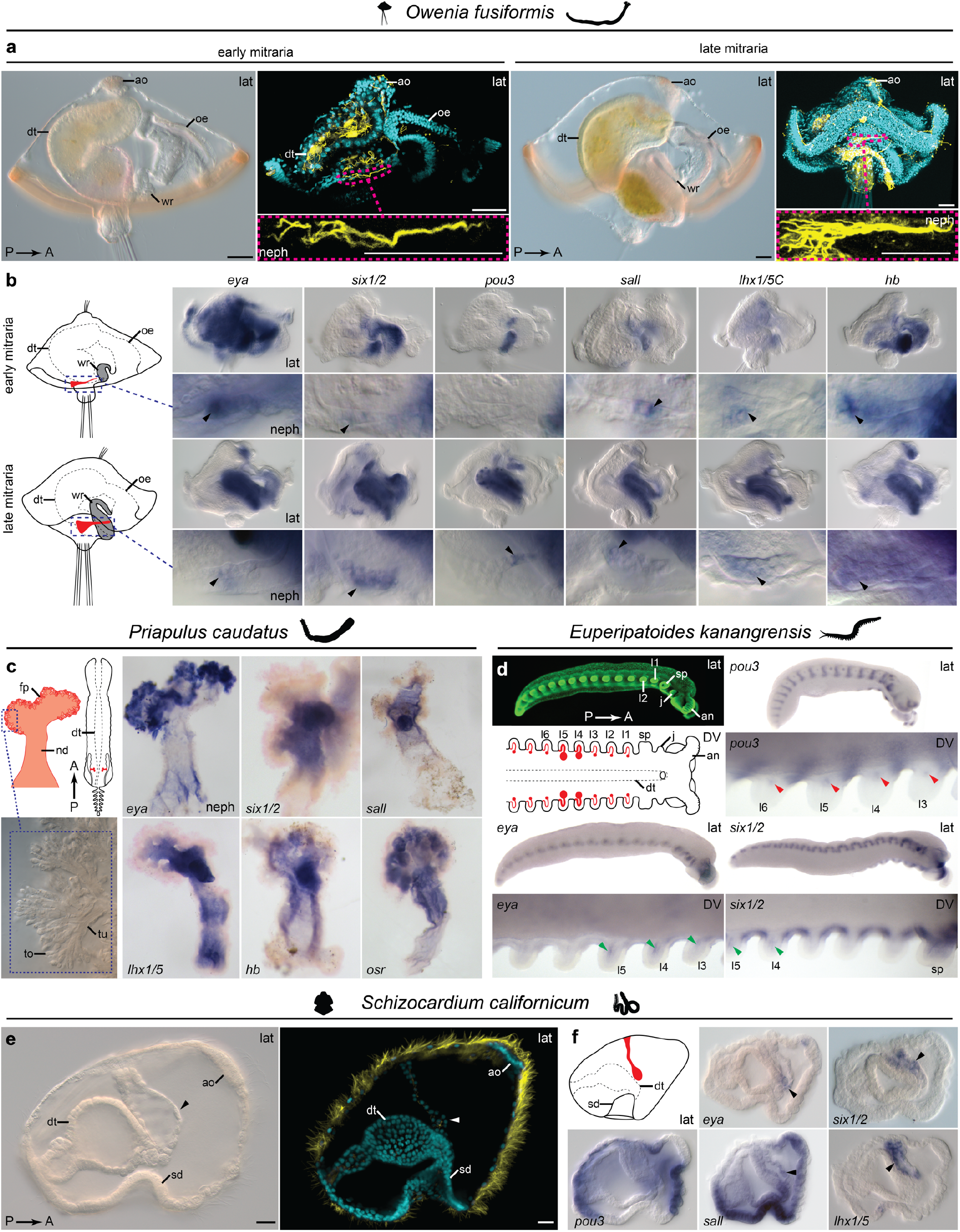
Expression of the nephridia-related transcription factors in annelid, ecdysozoans and hemichordate. **a**, A pair of ciliated protonephridia (outlined with magenta dotted line) is positioned on the lateral edges of the larval *O. fusiformis.* **b**, Transcription factors expressed in the developing protonephridia (black arrowheads) of *O. fusiformis.* **c**, Position and structure of the protonephridia of the adult *P. caudatus* and expression of the genes in the dissected organs. **d**, Embryo of *E. kanangrensis* possesses single metanephridium rudiment associated with each walking appendage, *pou3* is expressed in the proximal sacculi (red arrowheads), while *eya* and *six1/2* in the more distal canals (green arrowheads) of the developing organs. **e**, Early larva of *S. californicum* possesses ciliated, anterior coelomic vesicle, which serves as protonephridium (arrowheads). **f**, Expression of *eya*, *six1/2, sall* and *lhx1/5* is detected in the larval protonephridium (arrowheads), while pou3 is broadly expressed in the ectoderm and its expression is not detected in the excretory organ. Abbreviations: A, anterior; fa, frontal appendage; ao, apical organ; dt, digestive tract; vv, ventral view; fp, filtering portion; j, jaw; l1–l6, walking legs 1–6; lat, lateral view; nd, nephroduct; neph, magnified nephridium; oe, oesophagus; P, posterior; sd, stomodaeum; sp slime papilla; to, terminal organ; tu, tubule; wr, worm rudiment. DAPI stained cell nuclei are in cyan, Cybr-green stained cell nuclei in green and acetylated tubulin immunoreactivity is in yellow. Nephridia on the schematic drawings are marked in red. Scale bars, 20 μm.

### Molecular development of UF-based organs in Ecdysozoa

Since we showed that the expression of the developmental nephridial genes is highly conserved among various spiralian species, we examined whether a similar conservation is present in members of the second major protostome group, the Ecdysozoa. Two of the well-studied invertebrate model systems – the nematode *Caenorhabditis elegans* and the fly *Drosophila melanogaster* – belong to Ecdysozoa (Fig. 1a); however, both lack UF-based excretory organs and instead use presumably derived secretory excretory systems(3, 6) that develop without expressing the aforementioned conserved set of nephridial TFs(29, 30). Therefore, in order to reconstruct the ancestral molecular patterning of nephridiogenesis of the Ecdysozoa, it is essential to study species with UF-based excretory organs. Here, we investigated the expression of these conserved genes in two species that deploy UF for excretion, the priapulid *Priapulus caudatus* and the onychophoran *Euperipatoides kanangrensis.*

In the case of *P. caudatus,* we studied gene expression in the dissected adult protonephridia, which are part of the posteriorly positioned urogenital system (Fig. 4c)(31). In situ hybridization of *eya, six1/2, sall, lhx1/5, hb* and *osr* in the adult protonephridium (Fig. 4c) show that these genes are expressed in different portions of the organ – e.g. *eya* is mainly expressed in the terminal filtering portion, while expression of *six1/2*, *hb* and *lhx1/5* is restricted to the nephroduct (Fig. 4c).

Onychophorans possess serial, ciliated metanephridia in each trunk segment, which develop directly from the mesodermal rudiment at the base of each leg and, unlike the metanephridia of spiralians, are not preceded by any larval organs(32). Expression of some of the candidate nephridial genes has been already investigated in onychophorans and showed that *osr* and *lhx1/5* are indeed expressed in the metanephridial rudiments(33, 34), while *hunchback* does not seem to be directly involved in nephridiogenesis(35) (however it is expressed in the limb mesoderm, which might include some progenitor cells giving rise to the metanephridia(32)). Three of the remaining TFs (*eya*, *six1/2* and *pou3*) are expressed in the developing nephridia associated with the walking appendages (Figs. 4d, S4b). In the advanced embryos, *pou3* is expressed more proximally, likely in the filtering portion of the nephridium (the so-called sacculus), while *eya* and *six1/2* mark the more distal, nephridial canal (Fig. 4d). Signal from the probes against each gene is the strongest in the leg segments 4 and 5 (Figs. 4d, S4b), which corresponds to the presence of larger nephridia in those segments(32).

These results show that the conserved set of TFs expressed during the development of spiralian proto- and metanephridia is also likely to be involved in the formation and maintenance of corresponding ecdysozoan UF-based excretory organs, suggesting that these organs are homologues across all protostomes.

### Molecular development of nephridia in non-vertebrate deuterostomes

The genetic control of vertebrate kidney development is well described, however vertebrate kidneys represent evolutionarily derived excretory systems(6). To investigate gene expression during the development of deuterostomes with less derived excretory organs, we tested the expression of the conserved TFs in the indirect developing hemichordate *Schizocardium californicum*. Excretion in hemichordates is performed through an anterior coelomic compartment that initially functions as a protonephridium in the larva, and later gives rise to the adult metanephridial system with a podocyte-lined ultrafiltration site(36). In *S. californicum* the larval protonephridium appears in early larvae as an unpaired anterior coelomic vesicle connected by a ciliated canal with the dorsal epidermis (Fig. 4e)(37). *Eya*, *six1/2, sall* and *lhx1/5* are expressed in the larval protonephridium (Fig. 4f), with *eya* and *six1/2* restricted to the terminal and canal portion of the organ, respectively (Fig. 4f). Expression of *six1/2* and *eya* has been also reported in the presumptive larval nephridium of a sea urchin(38), another nonvertebrate deuterostome, which belongs to the sister group of hemichordates, the echinoderms. Furthermore, the expression of *lhx1/5* has been reported in the larval nephridium of amphioxus (39), supporting conservation of this TF also in the nephridial development of non-vertebrate chordates. We did not detect transcripts of *pou3* and *osr* in the developing protonephridium of the investigated life-stage of *S. californicum*. Instead, *pou3* shows a broad ectodermal expression (Fig. 4f), while *osr* is expressed in the digestive tract in the pattern similar to the one observed in lophophorates and annelids (Fig. 3d).

These results show that the molecular similarities of kidney and protostome nephridial development are also observed in the UF-based excretory system of non-vertebrate deuterostomes. Moreover, these data suggest that kidneys, proto- and metanephridia are homologous and that the conserved set of nephridial developmental TFs has been inherited from the last common nephrozoan ancestor.

### Conservation of the nephrozoan ultrafiltration proteins

We showed that the molecular similarities seen in the developing anatomically diverse UF-based excretory organs of Nephrozoa strongly suggest that those organs share an evolutionary ancestry. Next, we wanted to test whether these diverse excretory organs share the expression of genes encoding for structural proteins known for the formation of the extracellular filter in the excretory organs of vertebrates and flatworms(6, 9, 12). Such molecular conservation would further support the presence of a filtration apparatus necessary for UF and production of primary urine in the nephridia of the last common nephrozoan ancestor.

We investigated the expression of three structural genes (*nephrin, kirrel* and *ZO1*) in the phoronid, brachiopod, annelid, onychophoran and hemichordate. In phoronid, annelid and hemichordate, all three genes are expressed in the filtering cells of the excretory organs (Fig. 5a). In Onychophora, *nephrin* and *ZO1* are specifically expressed in the filtering sacculus at the base of each walking leg (Figs. 4a, S4c), however the *kirrel* homologue is only transiently expressed in the posterior portion of the embryo (Fig. S4c). In brachiopods, we did not observe expression of the UF genes in the larval nephridia (Fig. S2c), which are likely non-functional rudiments. In post-metamorphic juvenile brachiopods, the UF genes are broadly expressed (Fig. S2d), including expression in the periesophageal coelom, the presumptive site of ultrafiltration in adult brachiopods(40).

**Figure 5.**
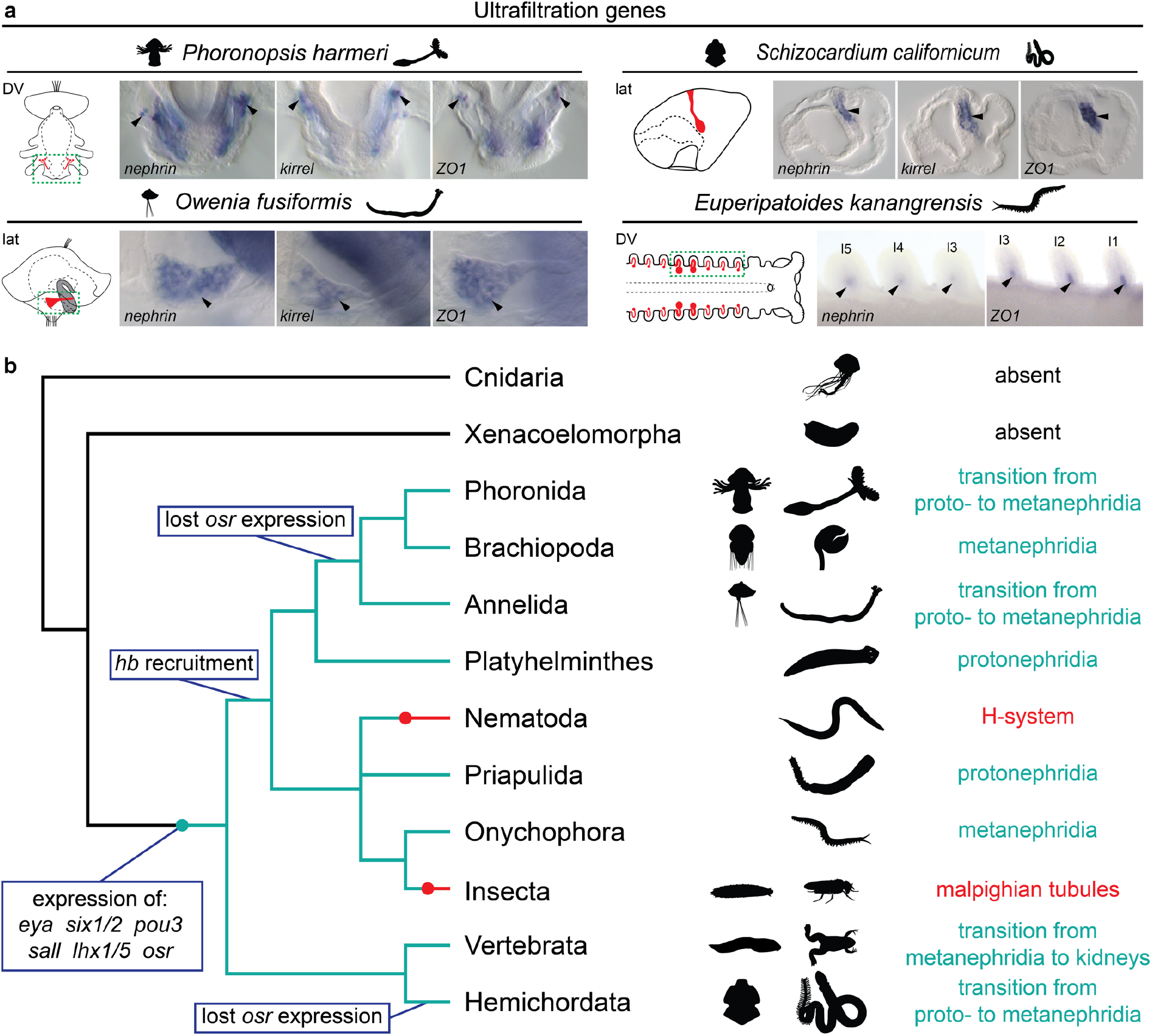
Conservation of the UF-related structural genes and evolution of the excretory organs in Nephrozoa. **a**, Putative UF-related structural genes are expressed in the filtering cells (arrowheads) of the excretory organs in various nephrozoan species. Areas shown on micrographs are outlined in green. **b**, Proposed scenario for the evolution of excretory organs ad their genetic control. The last common ancestor of Nephrozoa had ciliated, UF-based organs (light green), which were developing with an expression of the set of transcription factors, conserved in the contemporary members of the clade. The UF-based excretory organs has been lost and replaced by the secretion-based systems (red) in nematodes and insects. Abbreviations: vv, ventral view; l1–l5, walking legs 1–5; lat, lateral orientation. Nephridia on the schematic drawings are marked in red.

The observed conservation of the UF-genes among distantly related nephrozoans with ciliated excretory organs suggest that the last common nephrozoan ancestor already used a filter composed out of Nephrin, KIRREL and ZO1 proteins and deployed cilia-driven ultrafiltration as a mechanism for excretion. The orthologues of the UF-related genes are also present in the potential sister group of Nephrozoa, the xenacoelomorphs, where they are broadly expressed in tissues not related with the excretion process, e.g. in gonads and nervous system(41). This suggests that recruitment of those three genes into formation of excretory filter was an important step in the evolution of specialized excretory organs in Nephrozoa.

## Conclusions

We demonstrated that UF-based excretory organs in all major evolutionary lineages of Nephrozoa express a conserved set of transcription factors (*eya*, *six1/2, pou3, sall, lhx1/5, osr*) during their development, which indicates the presence of a deeply conserved nephrozoan molecular signature of the excretory organs (Fig. 5b). The observed lack of expression of some of those genes during development of excretory organs in particular species may be due to lineage specific losses (Fig. 5b). We propose that the observed conservation of molecular patterning is a result of the homology of all types of the nephrozoan UF-based excretory organs (kidneys, protonephridia and metanephridia), since they not only share molecular similarities but also show a morphological continuity of forms (e.g. ontogenetic transition from proto-to metanephridia(3, 11, 21, 22) or from metanephridia to kidneys(12)). Moreover, we provide evidence for the evolutionary conservation of the proteins involved in the formation of UF-apparatus, suggesting the presence of UF at the deepest nephrozoan node (Fig. 5b). The ancestral nature of UF is further supported by the distribution of UF-based organs on the phylogenetic tree (Fig. 1a) and the fact that the nephrozoan UF-sites share morphological similarities between distantly related clades(3, 6, 11–14). Therefore, despite lack of unambiguous fossil record of the nephrozoan ancestor, it is possible to reconstruct its excretory organs as ciliated and UF-based, likely similar to the simple protonephridia. This implies, that organs as divergent as protonephridia of microscopic worms that are composed out of just three cells(6, 42), and the enormous kidneys of the blue whale that weigh up to 360 kg(43) evolved from the same ancient organ present in the last common nephrozoan ancestor more than half a billion years ago.

Although we came closer to the understanding of the early evolution of excretory organs at the deep nephrozoan nodes, there are still many aspects of the evolution of excretion that require further investigation. In particular, the origin of excretory organs based on secretion, present in the most studied invertebrate model systems – nematodes (the so-called H-system) and insects (Malpighian tubules) – remains obscure. In *D. melanogaster* and *C. elegans* those organs neither express the nephrozoan ancient set of nephridial genes nor develop from any identifiable UF-based organs(29, 30, 44) and thus most likely emerged *de novo* (Fig. 5b). Besides Malpighian tubules, insects also possess specialized excretory cells – nephrocytes – that perform an ultrafiltration-like process with a filter composed of Nephrin, KIRREL and ZO1(45). The presence of this conserved UF apparatus as well as morphological data from crustaceans, in which similar nephrocytes develop from metanephridial podocytes(46), indicate that the UF organs of insects might become reduced to the single cell structures(13), in tandem with the development of Malpighian tubules. Further investigation of the molecular basis of the development of Malpighian tubules and H-system in additional non-model arthropod and nematode species, is needed in order to understand how ancestral nephrozoan excretory organs got replaced by entirely new systems in those two evolutionarily successful groups.

## Materials and methods

### Animal collections and sample fixations

Adult gravid animals were collected from Bodega Bay, California, USA (*P. harmerĩ),* San Juan Island, Washington, USA (*T. transversa),* Station Biologique de Roscoff, France (*O. fusiformis*), Kanangra Boyd National Park, NSW, Australia (*E. kanangrensis*) and Morro Bay State Park, California, USA (*S. californicum).* The animals were spawned and larvae or embryos were obtained as described elsewhere(26, 33, 37, 47, 48). Adult *P. caudatus* worms were collected from Gullmarsfjorden, Sweden and dissected in laboratory to obtain their protonephridia. Before fixation, larval were relaxed in 7.4% magnesium chloride. All samples were fixed in 4% paraformaldehyde for 1 h at room temperature. After fixation, samples were washed in 0.1% Tween 20 phosphate buffer saline, dehydrated through a graded series of methanol, and stored at −20 °C in pure methanol or ethanol.

### Gene identification

Transcriptomes from mixed developmental stages and/or adults were used for gene identification. Gene orthologs were identified based on reciprocal tBLASTn search and confirmed with phylogenetic analysis (Figs. S5 and 6). Protein sequences were aligned with reference sequences downloaded from GenBank (Supplementary Tables 1 – 7) using CLC Main Workbench 7 and non-conserved regions where removed with TrimAl (using the gappyout option). Phylogenetic analyses were performed in FastTree v2.1(49) (using the LG amino acid substitution model) and in RaxML(50) (using the best fitted model, chosen separately for each protein with ProtTest 3.4.2(51)).

### Immunohistochemistry and in situ hybridization

Morphology of studied stages was investigated with mouse primary monoclonal antibodies against acetylated tubulin (Sigma, T6793) in 1:500 concentration, visualized with the secondary goat anti-mouse antibodies (Life Technologies) conjugated with fluorochrome (AlexaFluor647) in 1:50 concentration. Cell nuclei were stained with DAPI (*P. harmeri, T. transversa, O. fusiformis, S. californicum*) or SYBR-green (*E. kanangrensis*) and samples were mounted in 80% glycerol (*P. harmeri, S. californicum*) or Murray Clear (*T. transversa, O. fusiformis*).

Single whole-mount colorimetric and fluorescent in situ hybridization was performed following an established protocols ((33)for *E. kanangrensis* and (52) for the other species) with probe concentration of 1 ng/μl and hybridization temperature of 67°C. Proteinase K treatment time was adjusted for each species and ranged from 2 (*P. harmeri, O. fusiformis, P. caudatus, S. californicum*) to 10 min (*T. transversa).* Samples were mounted in Murray Clear (fluorescently stained *T. transversa* larvae and nephridia of *P. caudatus*) or 70% glycerol (all remaining samples). Double fluorescent whole-mount in situ hybridization was performed as described elsewhere(53) and samples were mounted in 80% glycerol.

### Imaging and image processing

Fluorescently labeled samples (immunohistochemical staining and fluorescent in situ) were scanned in a Leica SP5 confocal laser-scanning microscope. Samples investigated with colorimetric in situ hybridization were imaged with Zeiss AxioCam HRc connected to a Zeiss Axioscope Ax10 compound scope with bright-field Nomarski optics or Zeiss AxioCam MRc connected to Zeiss Discovery.V8 dissecting scope. Images were analyzed and adjusted for brightness and contrast with IMARIS 9.1.2 (confocal scans) and Photoshop CS6 (Adobe) (light micrographs) and figure plates were assembled with Illustrator CS6 (Adobe). Some of the used silhouettes of animals were downloaded from PhyloPic.com.

## Data availability

All newly determined sequences have been deposited in GenBank under accession numbers MT900856–MT900925. Multiple protein alignments used for orthology assignments are available upon request from the corresponding author.

## Acknowledgements

We thank all members of the Hejnol lab for their continuous support and animal care and collections. We thank Karl Menard from Bodega Bay Marine Lab for help with collecting *P. harmeri* adults. We are grateful to the “Centennial” boat crew of Friday Harbor Marine Laboratories for *T. transversa* collections, the Espeland marine biological station personnel for the *N. anomala* collections, and the Sven Loven Centre and the “Oscar von Sydow” crew for the *P. caudatus* collections. We are further grateful to the Roscoff Marine Station for the collections of adult *O. fusiformis* and thankful for the support of the New South Wales Government Department of Environment and Climate Change by provision of a permit SL100159 to collect onychophorans at Kanangra-Boyd National Park. We thank Auston Rutledge, for support of the husbandry of *S. californicum*. This study was supported by the European Research Council (ERC) Grant Agreement No. 648861 to A.H. The collections of *P. caudatus* was funded by ASSEMBLE+ grants to A.H. and C.A., G.B. received support from the Vetenskåpsradet (Swedish Research Council) Grant number VR 2015-04726 and C.L. received support from NASA NNX13AI68G.

## Author contributions

A.H. conceived the study. L.G., C.A., A.H. and R.J. designed experiments and analyses. L.G., C.A., P.B., R. J. and A.H. performed collections. L.G. and R. J. performed phylogenetic analyses and gene family evolution studies. L.G., C.A., and R.J. performed gene expression analyses. All authors contributed to interpretation of the results, and L.G. and A.H. drafted the manuscript. C.A., C.J. and G.B. edited the manuscript.

