## Supplementary Information for "A single origin of animal excretory organs"

**This PDF file includes:**

Figures S1 to S7

Tables S1 to S7

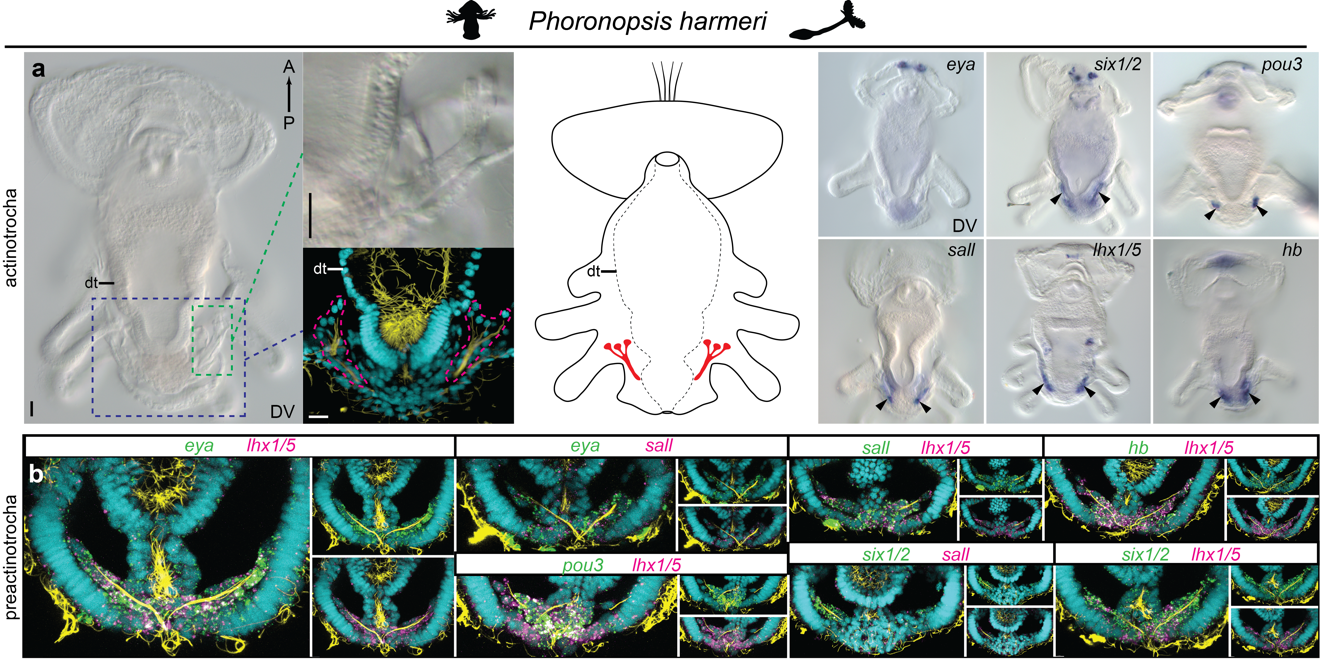

Fig. S1. Expression of the nephridia-related transcription factors in *P. harmeri*. a, Morphology of protonephridia and gene expression in the organs (arrowheads) of the advanced phoronid larva. Ciliated cells forming branching protonephridia are outlined in magenta. b, Co-expression of the investigated transcription factors in the protonephridia of preactinotrocha larva of *P. harmeri*. Abbreviations: A, anterior; dt, digestive tract; vv, ventral view; P, posterior. DAPI stained cell nuclei are in cyan and acetylated tubulin immunoreactivity is in yellow. Nephridia on the schematic drawing are marked in red. Scale bars, 20 μm.

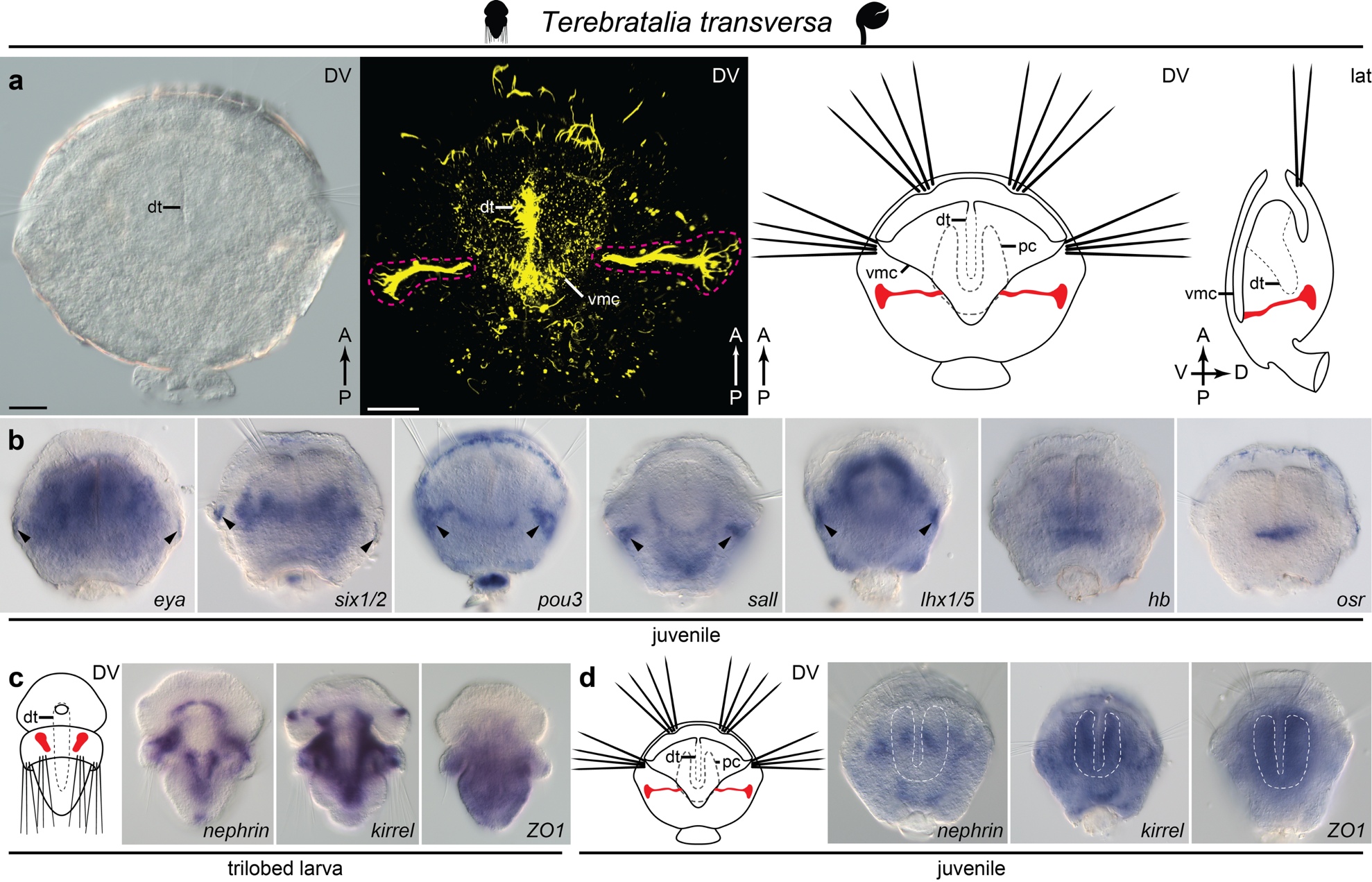

**Fig. S2.** Expression of the nephridia-related transcription factors and structural genes in *T. transversa*. **a**, A postmetamorphic juvenile of *T. transversa* possesses pair of ciliated metanephridia opening to the ventral mantle cavity. **b**, Transcription factors *eya*, *six1/2*, *pou3*, *sall* and *lhx1/5* are all expressed in the metanephridia (arrowheads) also after metamorphosis. **c**, Expression of the UF-related genes cannot be detected in the larval nephridial rudiments. **d**, The UF-related genes have broad expression domains in the postmetamorphic juveniles, which include cells in the periesophageal coelom (outlined in white), the putative UF-site of the adult brachiopods. Abbreviations: A, anterior; D, dorsal; dt, digestive tract; vv, ventral view; lat, lateral view; P, posterior; pc, periesophageal coelom; V, ventral; vmc, ventral mantle cavity. Acetylated tubulin immunoreactivity is in yellow. Nephridia on the schematic drawings are marked in red. Scale bars, 20 μm.

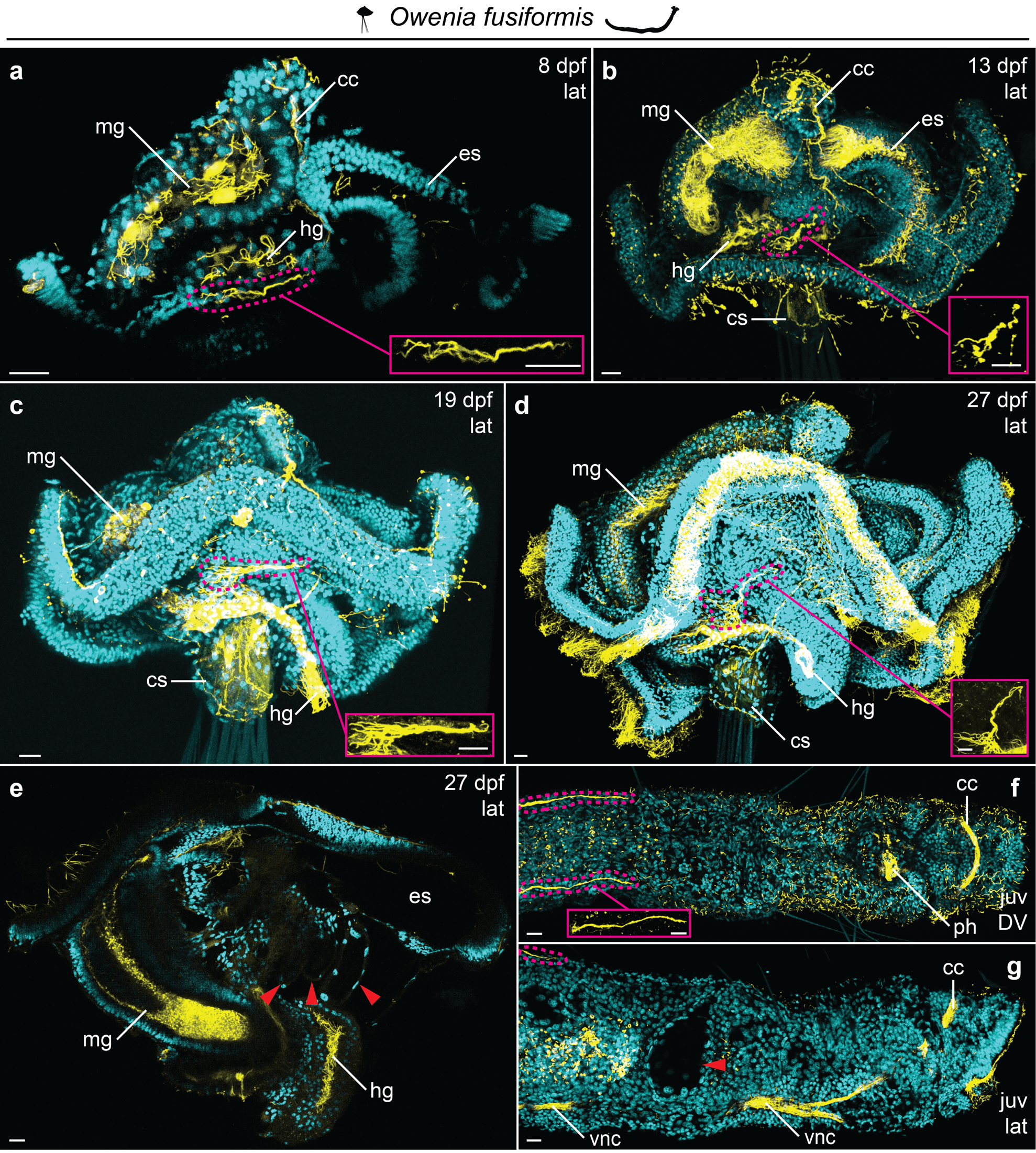

**Fig. S3**. Development of the excretory organs in *O. fusiformis*. The larval, ciliated protonephridium (outlined in magenta dotted lines) can be observed from 8 days post fertilization (dpf) (**a**), throughout later larval stages (**b**, **c**). In the competent larva (**d**) the nephridium is present and partially incorporated into the epidermis of the worm rudiment. **e**, The empty cavities, described as larval metanephridial rudiments, are not ciliated (red arrowheads) and likely they represent rudiments of the future tube-secreting glands. **f**, Immediately after metamorphosis a pair of ciliated rudiments of adult metanephridia is present on the dorsal side of the juvenile worm. They have similar size and shape to the nephridia of competent larvae. **g**, The empty cavities, described before as juvenile metanephridia (red arrowhead), are not ciliated, we suggest that they represent tube-secreting glands. The nephridia are instead positioned on the dorsal side (outlined in magenta dotted line), posteriorly to the first tube-secreting gland. Abbreviations: cc, circumesophageal connective; cs, chaetal sac; vv, ventral view; es, esophagus; hg, hindgut; juv, juvenile; lat, lateral view; mg, midgut; ph, pharynx; vnc, ventral nerve cord. DAPI stained cell nuclei are in cyan and acetylated tubulin immunoreactivity is in yellow. Scale bars, 10 μm.

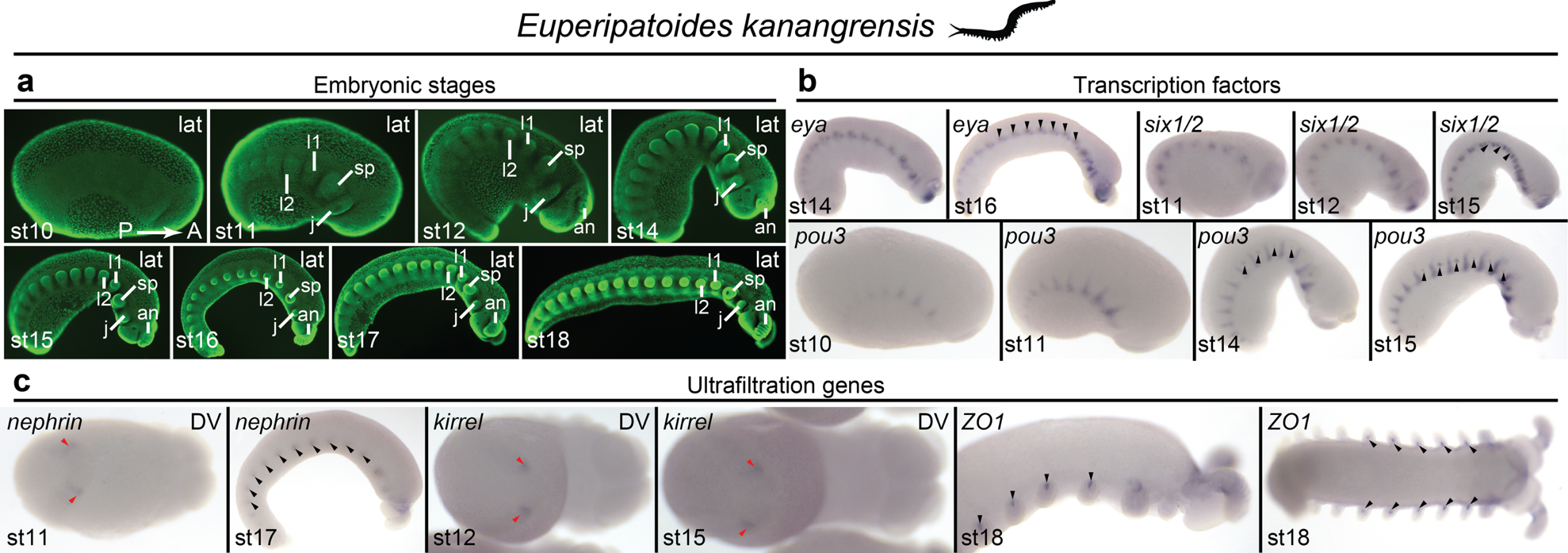

**Fig. S4.** Expression of the nephridia-related transcription factors and structural genes in *E. kanangrensis*. **a**, Staging and morphology of the developing onychophoran embryo, Cybr-green stained cell nuclei in green. **b**, Expression of the nephridia-related transcription factors in the metanephridial rudiments (black arrowheads). Onset of the nephridial expression of each gene is initiated at the different embryonic stage. **c**, Expression of the putative UF-related genes: *nephrin* and *ZO1* are expressed in the sacculi of the developing metanephridia (black arrowheads), while *kirrel* is transiently expressed in a pair of posterior domains (red arrowheads), which also express nephrin at the embryonic stage 11. Abbreviations: A, anterior; fa, frontal appendage; vv, ventral view; j, jaws; l1–l2, walking legs 1–2; lat, lateral view; P, posterior; sp, slime papillae; st10–st18, embryonic stages 10–18.

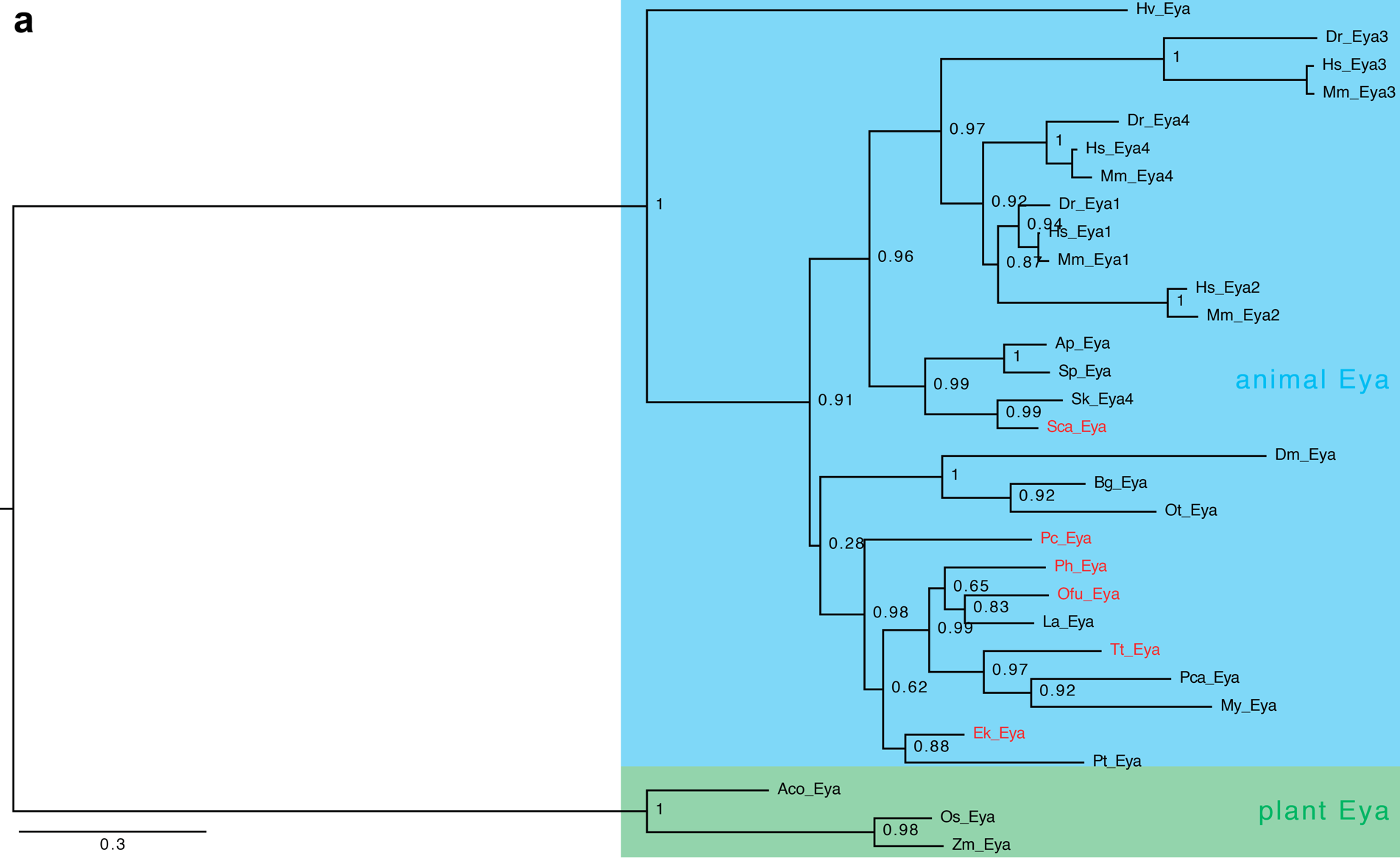

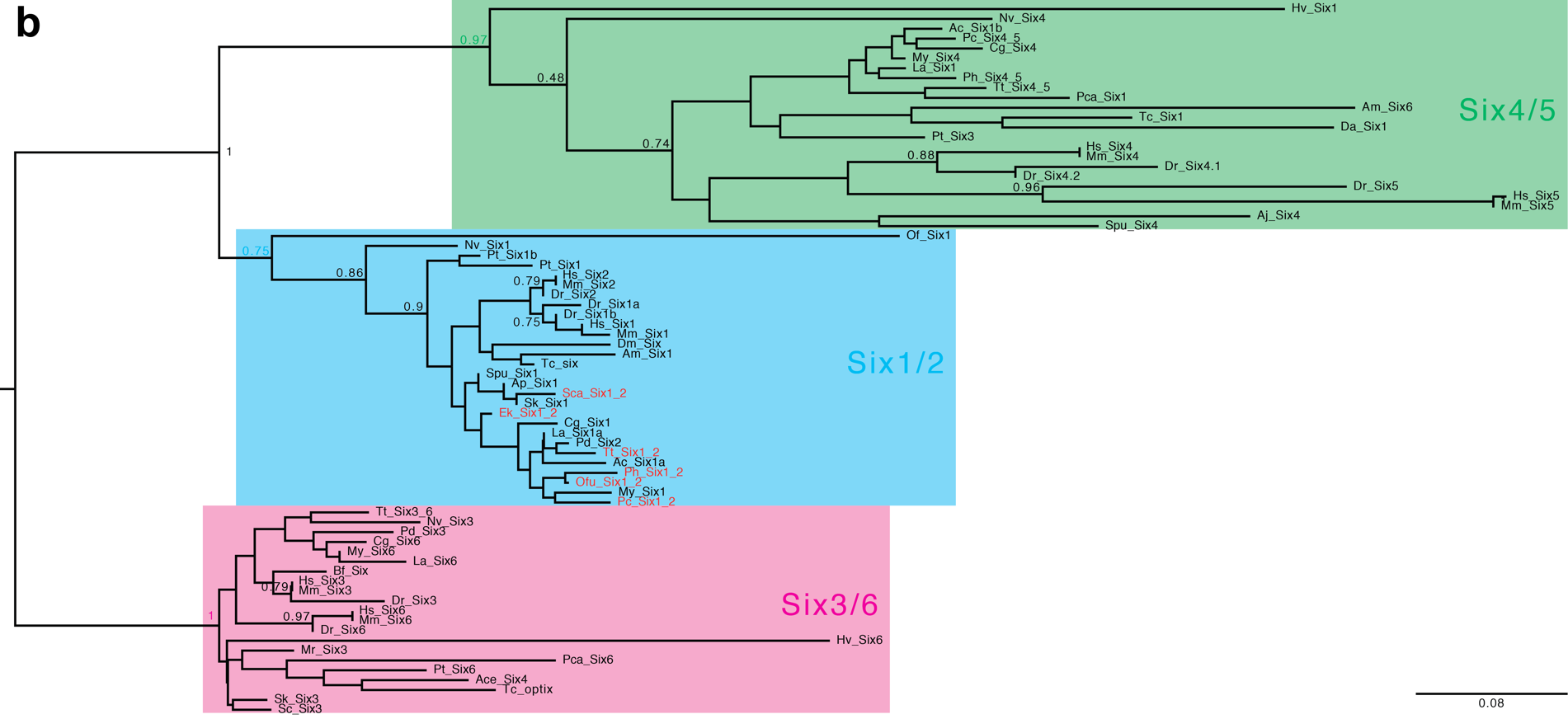

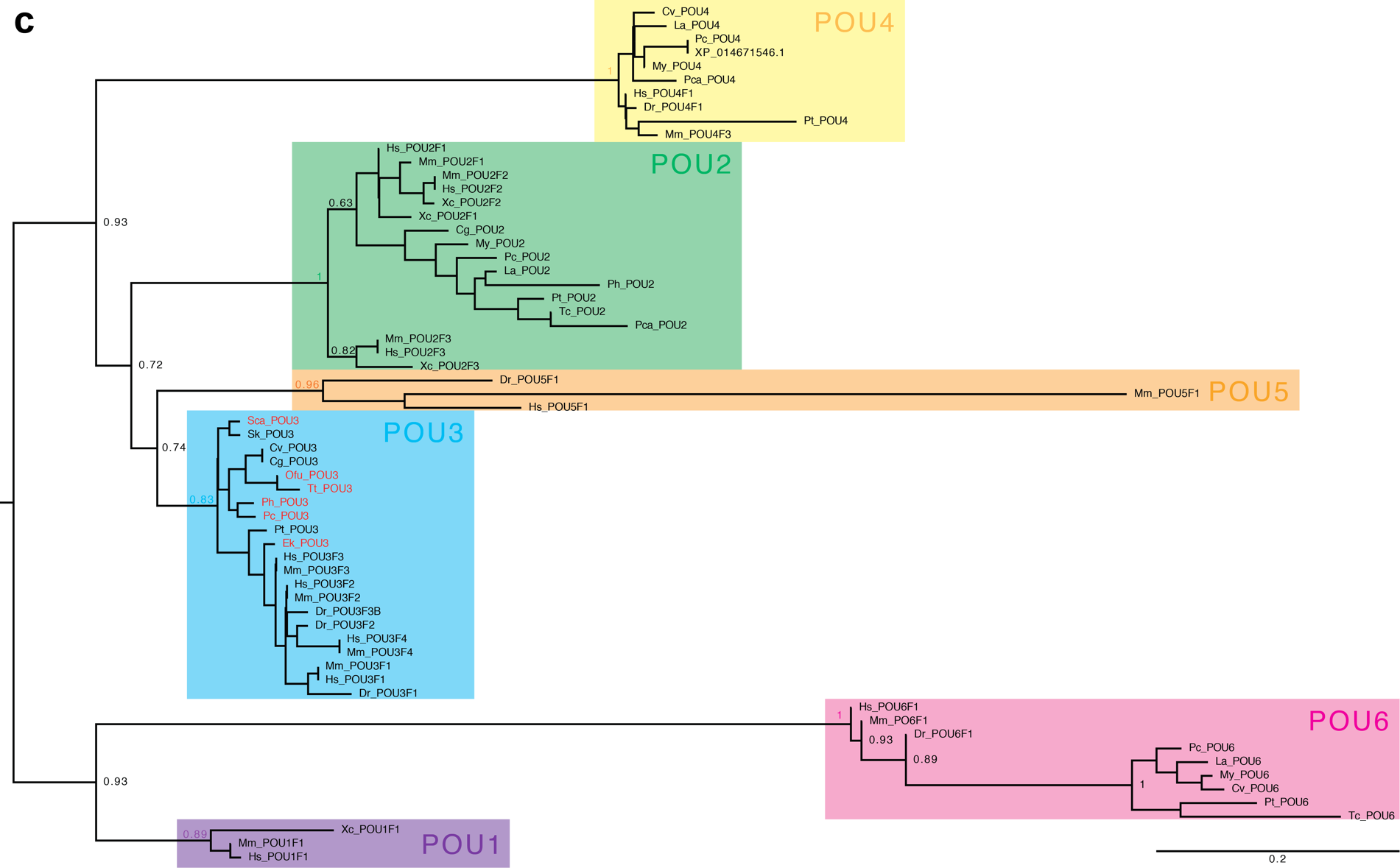

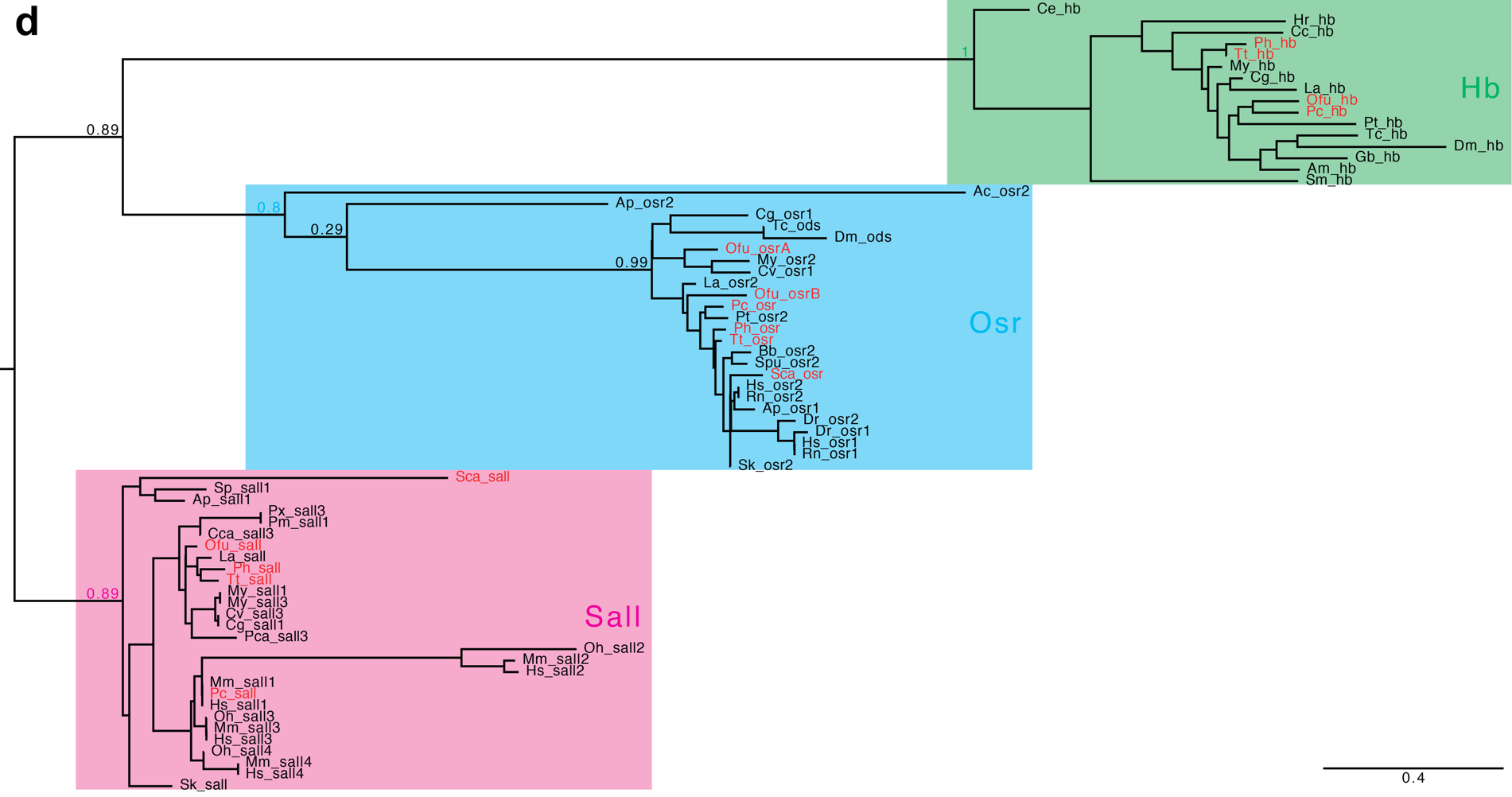

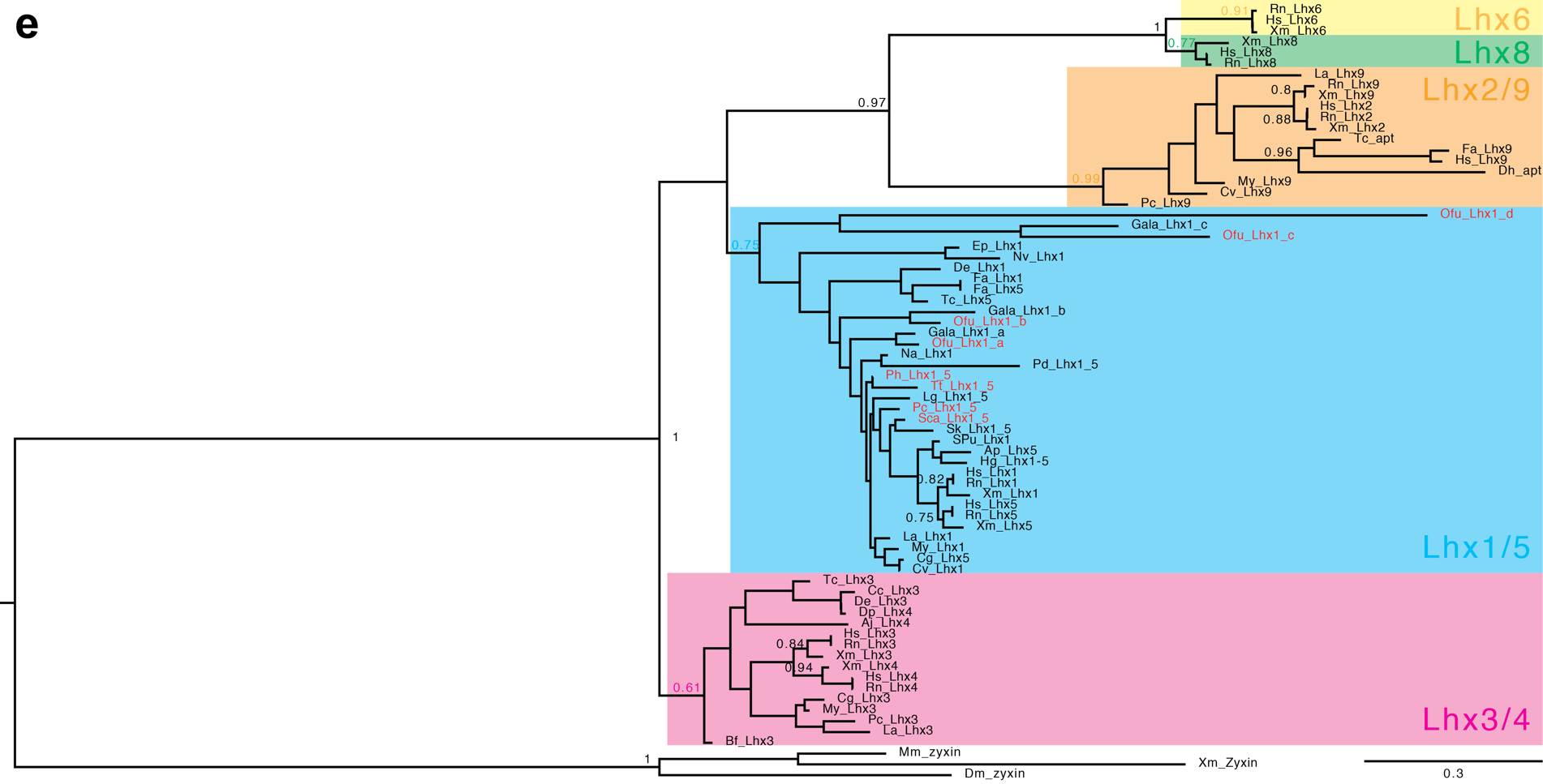

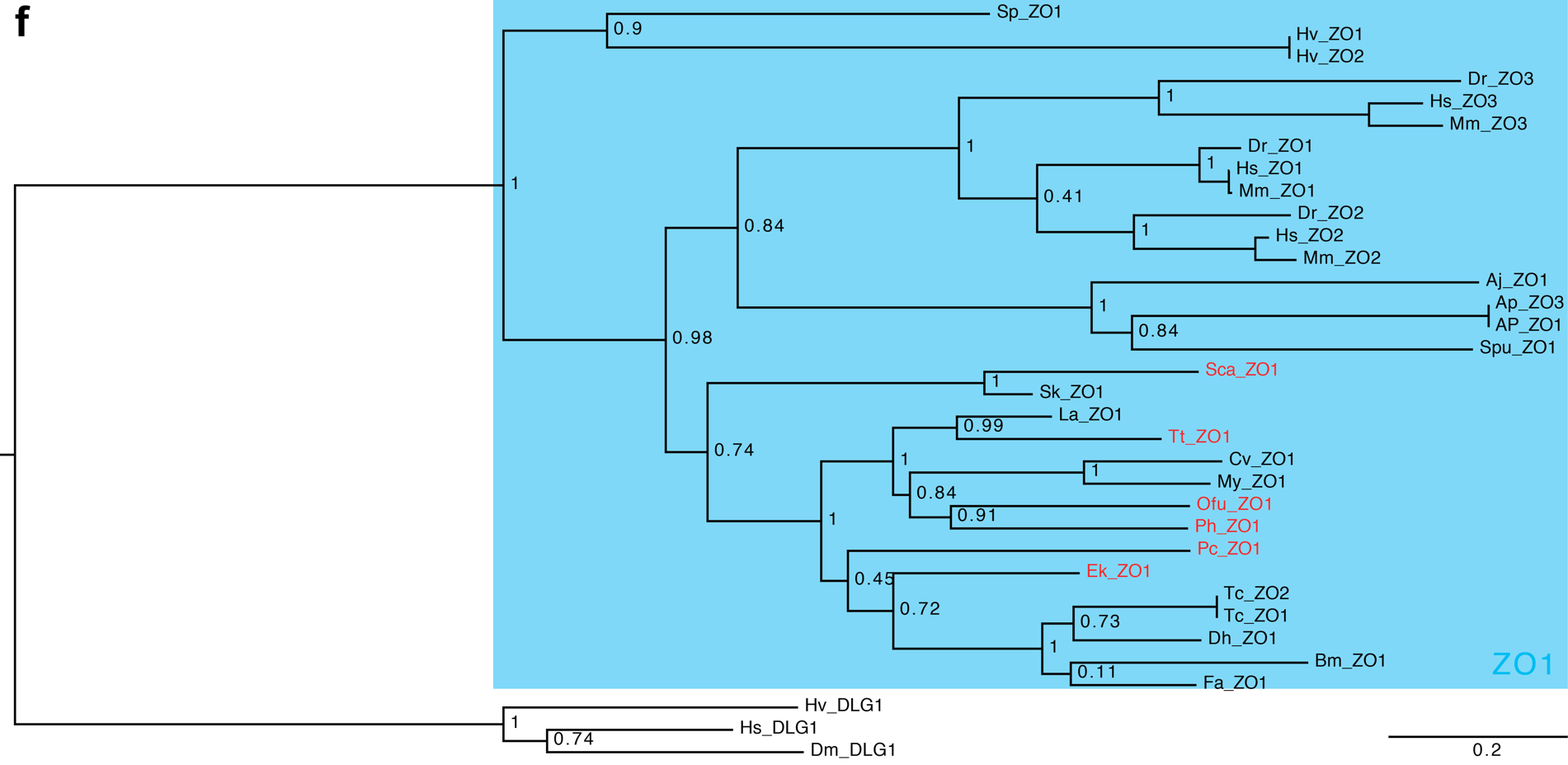

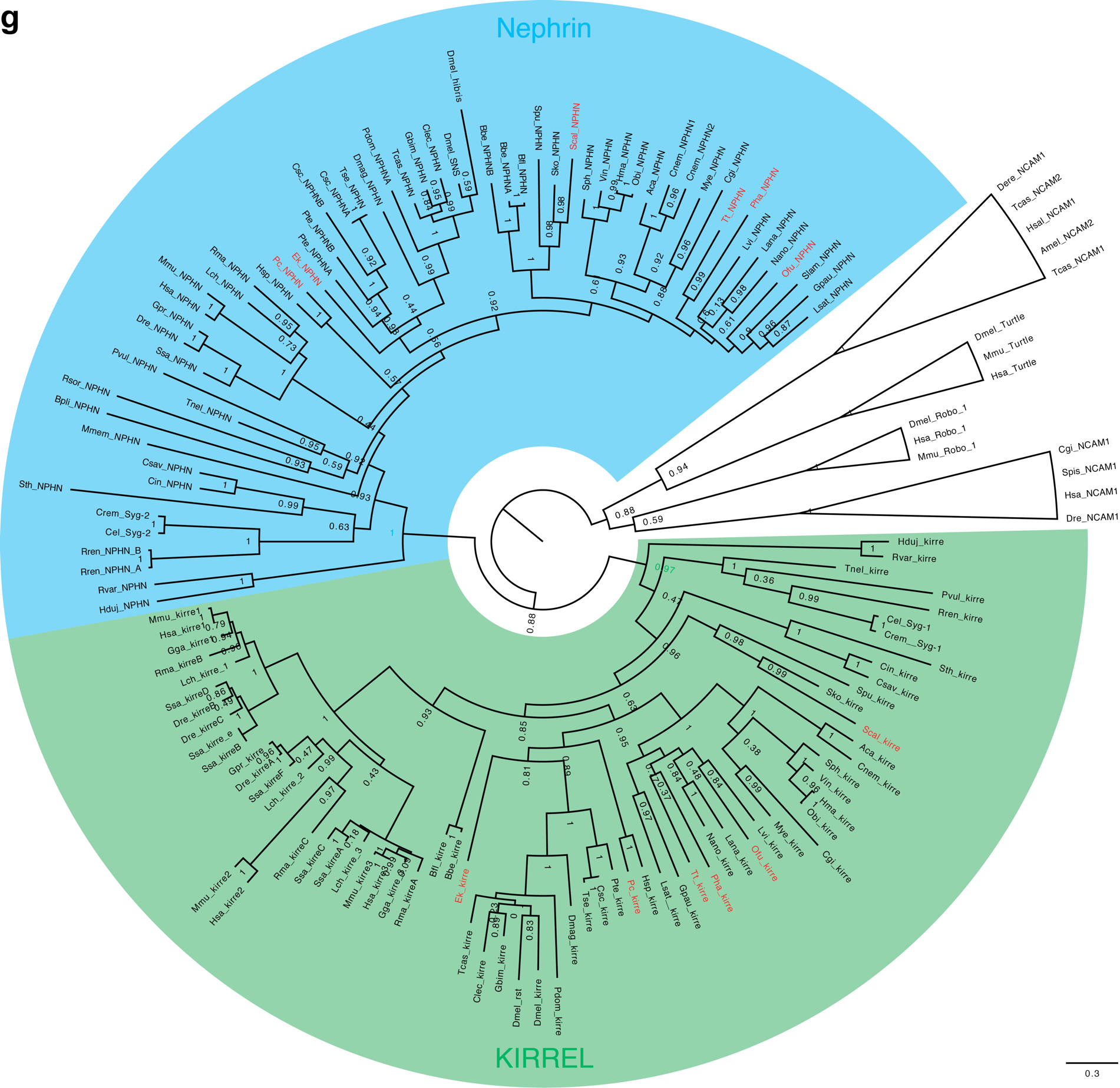

**Fig. S5.** Orthology analyses (**a**–**g**). Trees were generated with FastTree software under LG amino acid substitution model. SH-like support values are shown for the important nodes. Genes from studied species are marked in red. Full species names and sequences accession number are provided in Supplementary tables 1–7.

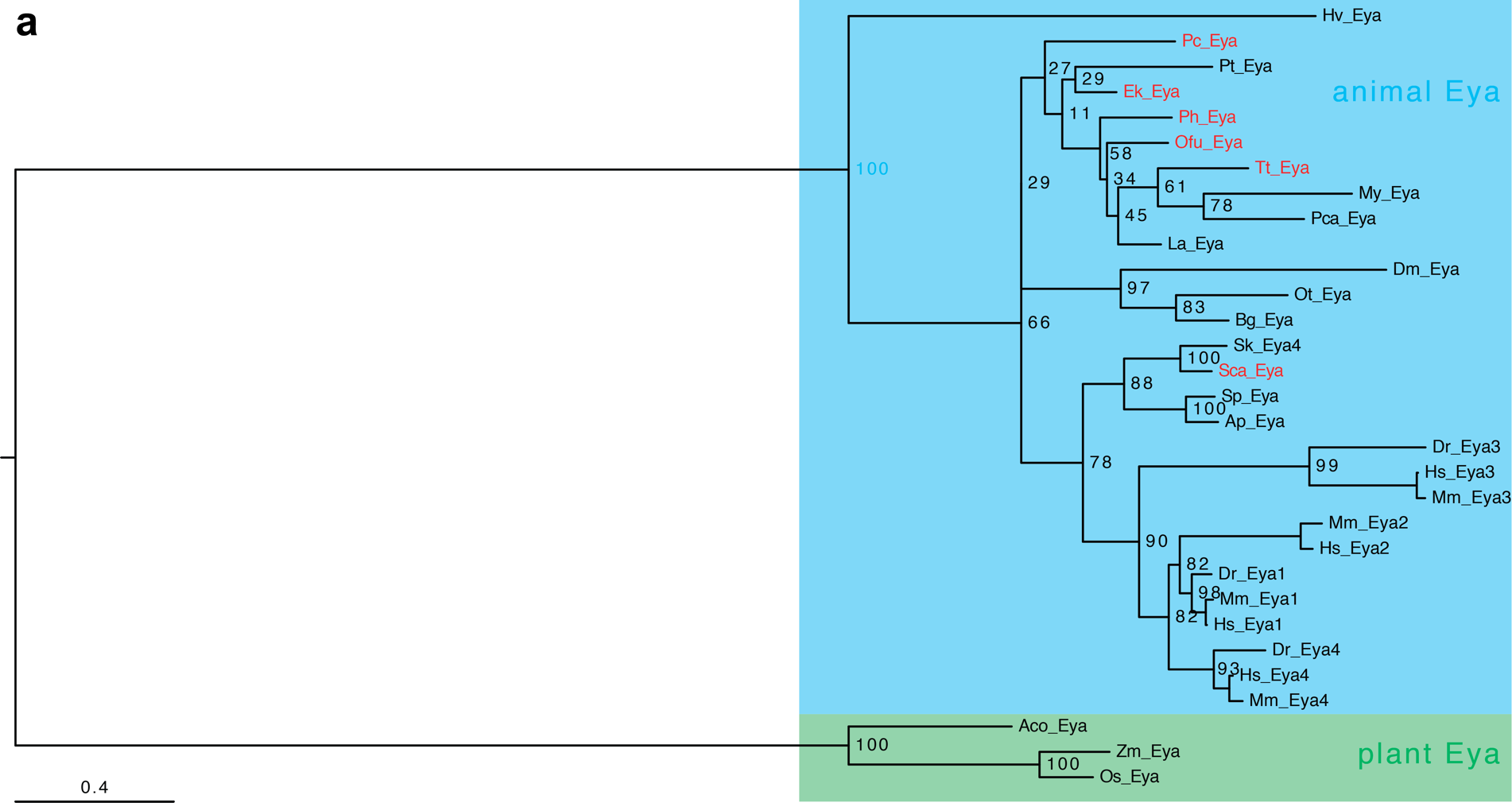

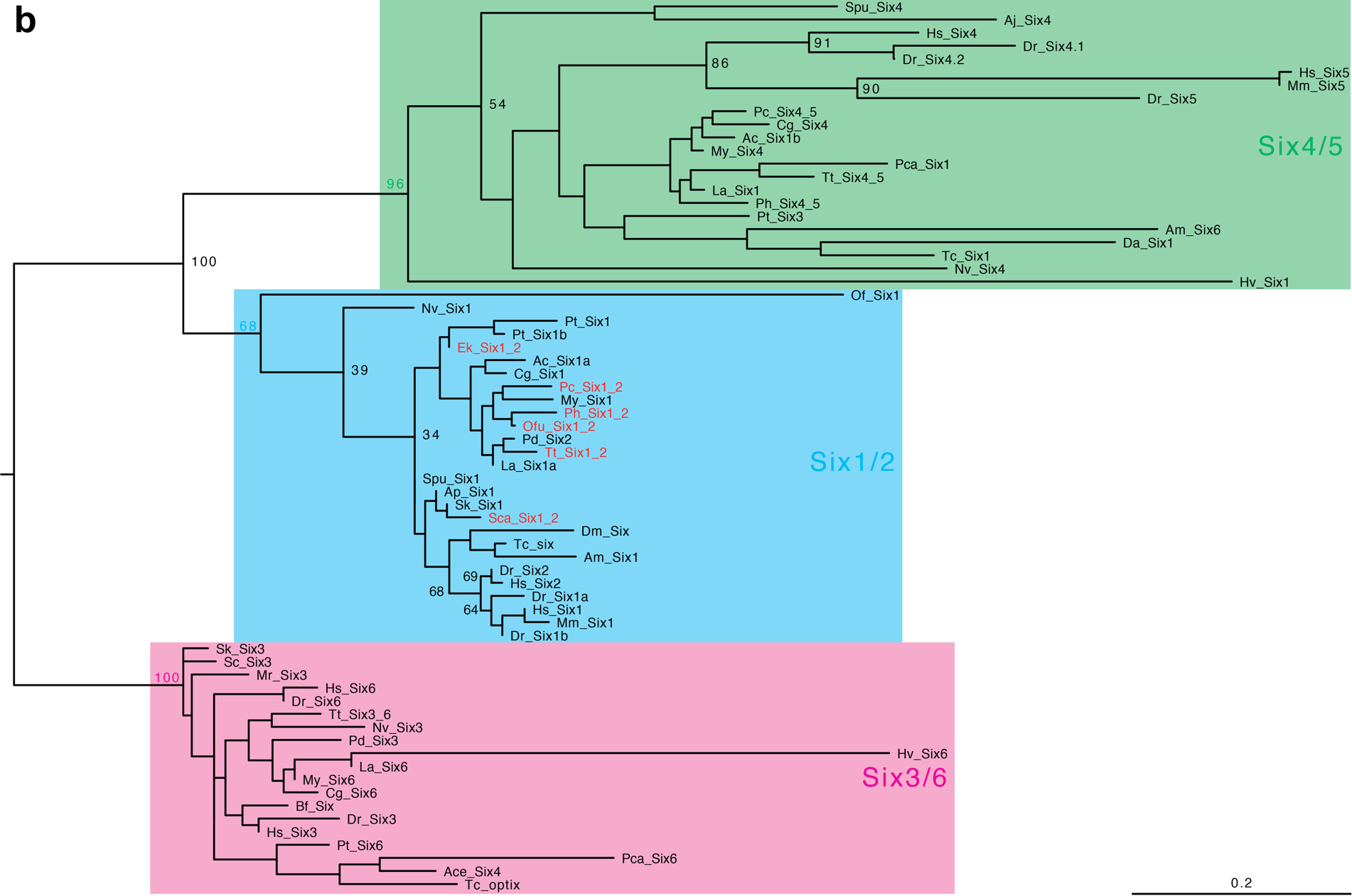

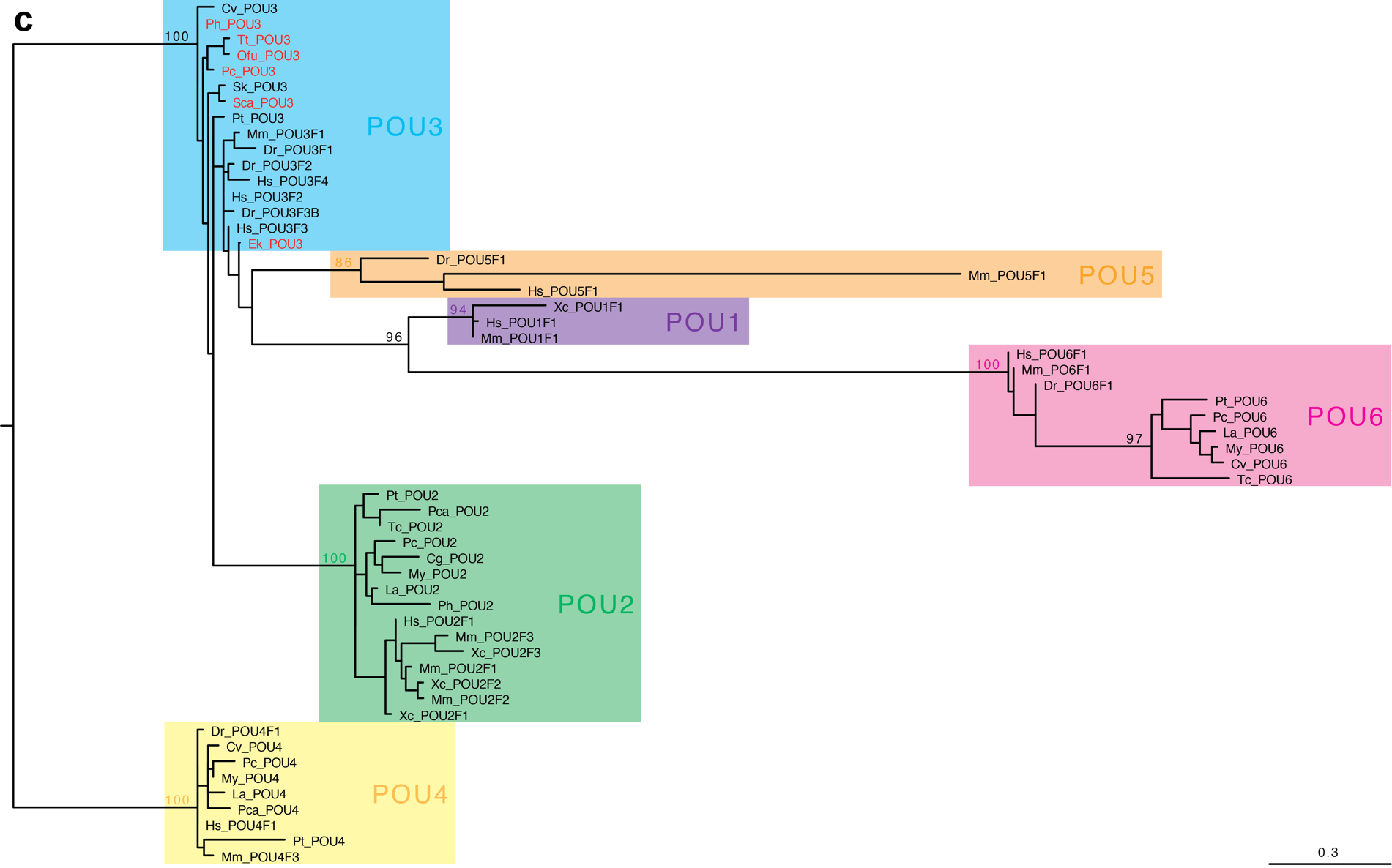

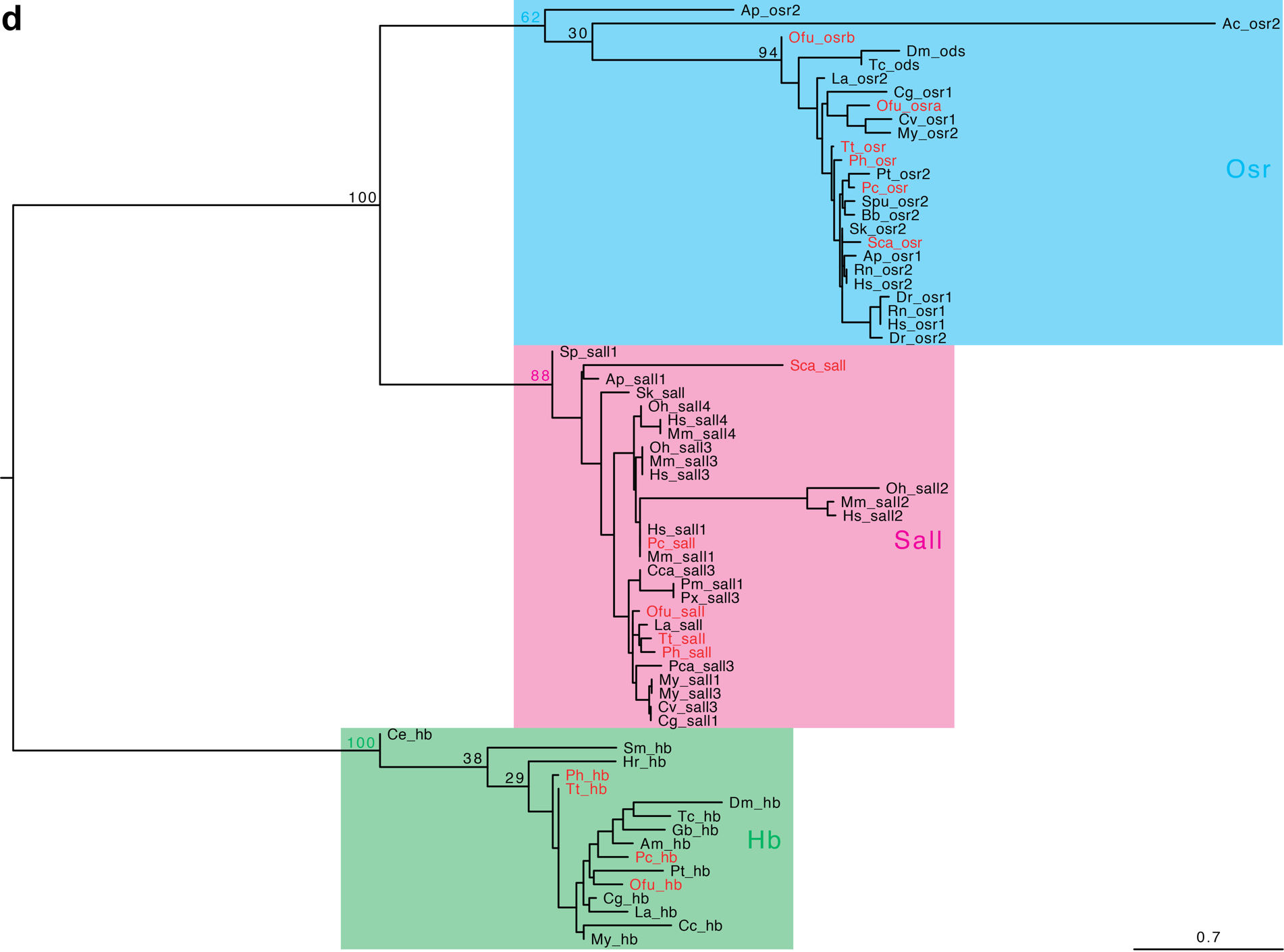

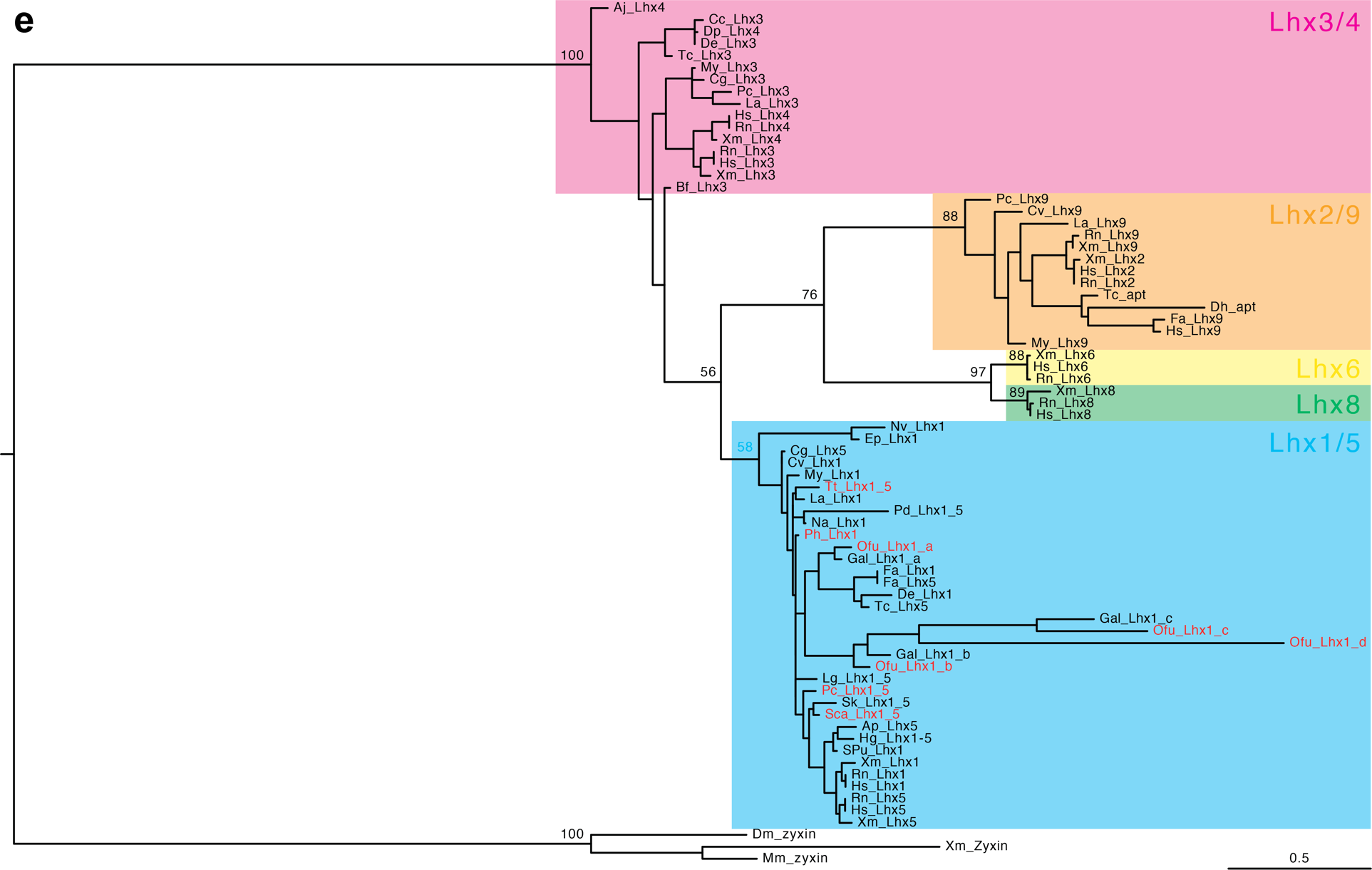

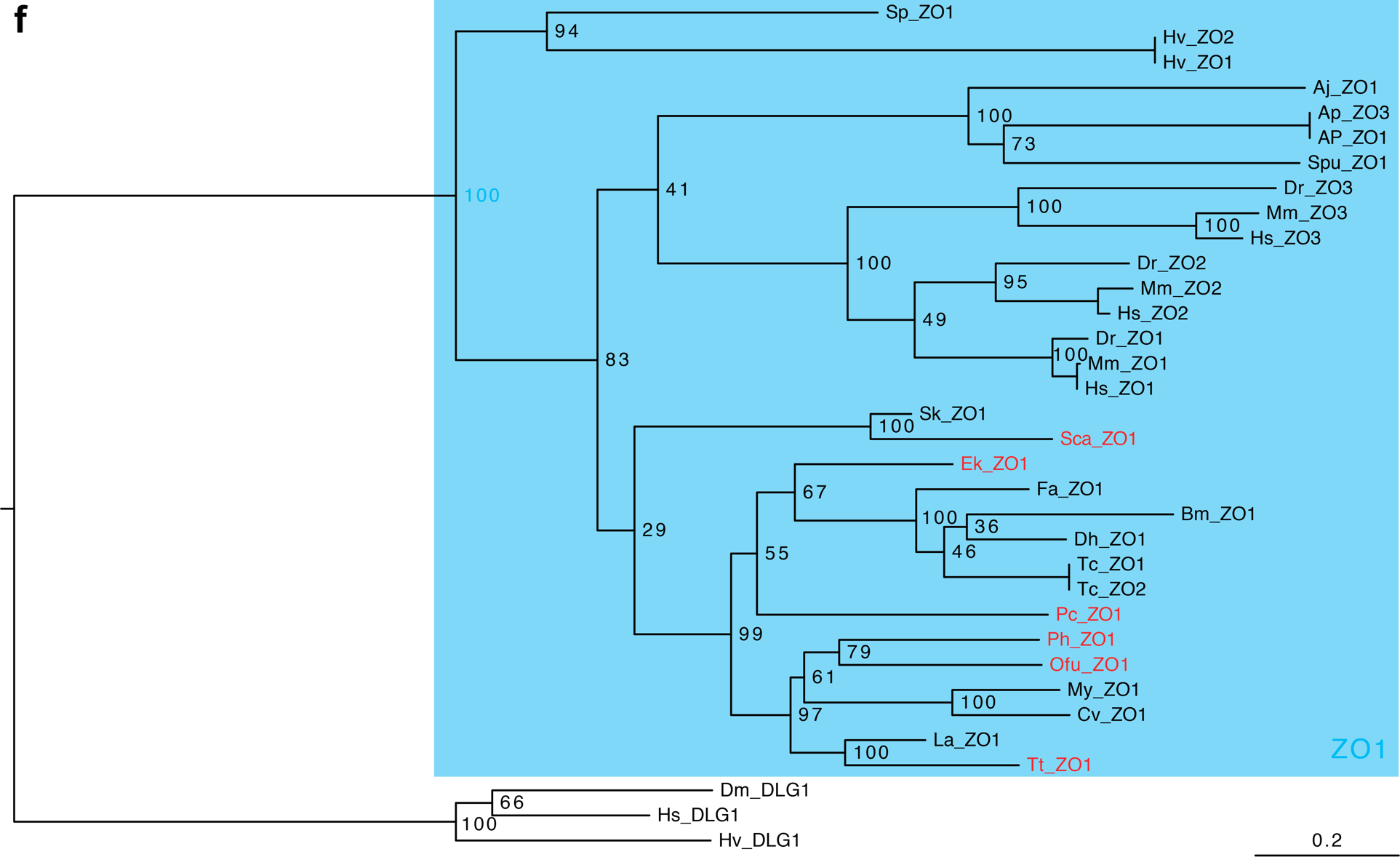

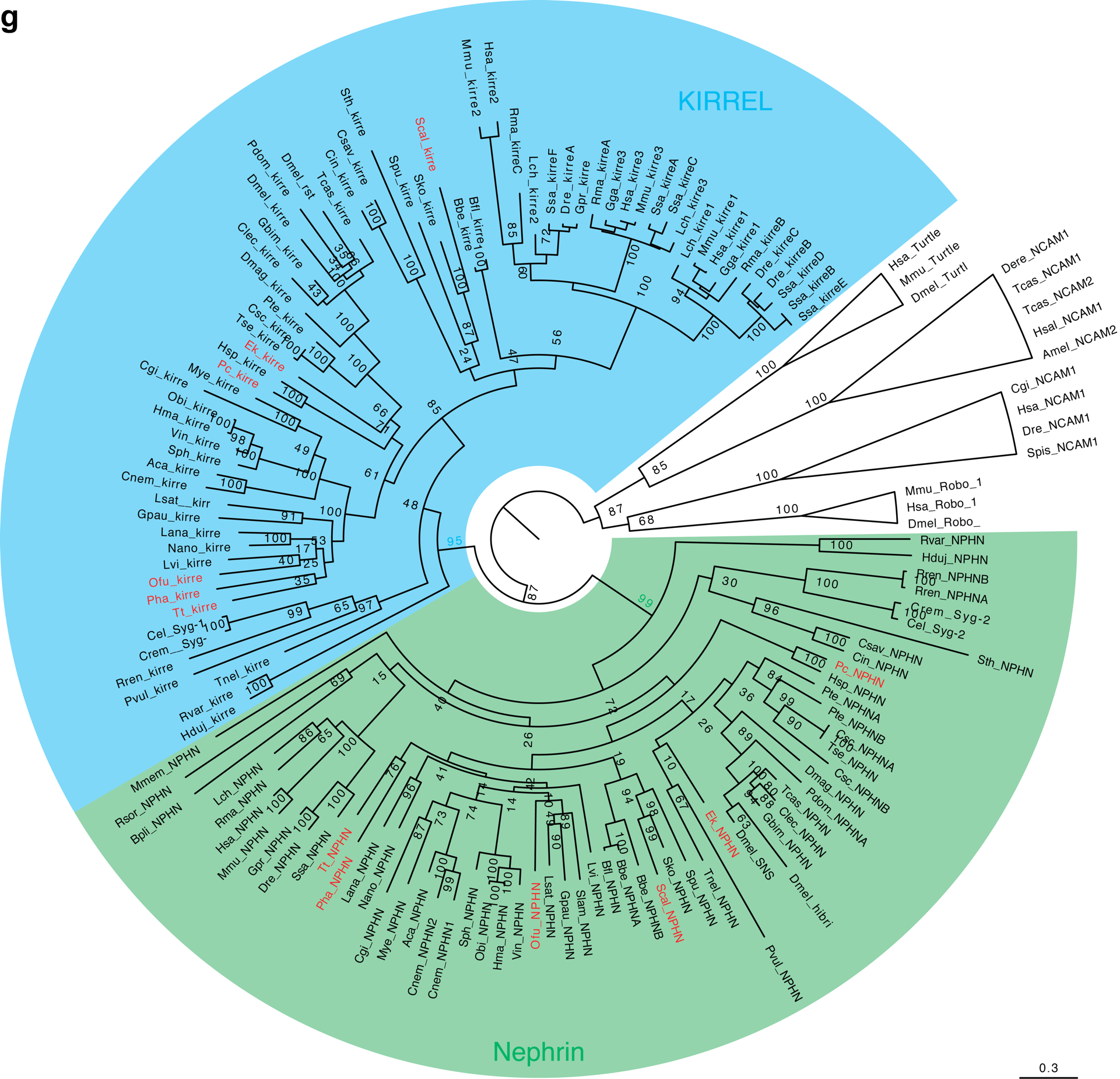

**Fig. S6.** Orthology analyses (a–g). Maximum likelihood trees were generated with RaxML software under JTT+I+G+F (a), LG+I+G (b, f, g), LG+G (c, d) and JTT+I+G (e) amino acid substitution models. Bootstrap values are shown for the important nodes. Genes from studied species are marked in red. Full species names and sequences accession number are provided in Supplementary tables 1–7.

**Table S1** Sequences used for the assessment of *eya* genes orthology

| **tree label** | **protein** | **species** | **clade** | **source** | **sequence** |
| --- | --- | --- | --- | --- | --- |
| Dr_Eya4 | eyes absent homolog 4 | *Danio rerio* | Vertebrata | protein from NCBI | XP_021325474.1 |
| Hs_Eya4 | eyes absent homolog 4 | *Homo sapiens* | Vertebrata | protein from NCBI | NP_742101.2 |
| Mm_Eya4 | eyes absent homolog 4 | *Mus musculus* | Vertebrata | protein from NCBI | XP_017169287.1 |
| Sk_Eya | eyes absent homolog 4 | *Saccoglossus kowalevskii* | Hemichordata | protein from NCBI | XP_006817779.1 |
| Dr_Eya3 | eyes absent homolog 3 | *Danio rerio* | Vertebrata | protein from NCBI | XP_005170122.1 |
| Hs_Eya3 | eyes absent homolog 3 | *Homo sapiens* | Vertebrata | protein from NCBI | NP_001981.2 |
| Mm_Eya3 | eyes absent homolog 3 | *Mus musculus* | Vertebrata | protein from NCBI | XP_006538600.1 |
| Hs_Eya2 | eyes absent homolog 2 | *Homo sapiens* | Vertebrata | protein from NCBI | NP_005235.3 |
| Mm_Eya2 | eyes absent homolog 2 | *Mus musculus* | Vertebrata | protein from NCBI | NP_001258892.1 |
| Ap_Eya | eyes absent homolog 1 | *Acanthaster planci* | Echinodermata | protein from NCBI | XP_022089633.1 |
| Bg_Eya | eyes absent homolog 1 | *Blattella germanica* | Arthropoda | protein from NCBI | PSN54252.1 |
| Dr_Eya1 | eyes absent homolog 1 | *Danio rerio* | Vertebrata | protein from NCBI | XP_009295602.1 |
| Dm_Eya | eyes absent homolog 1 | *Drosophila melanogaster* | Arthropoda | protein from NCBI | NP_723188.1 |
| Hs_Eya1 | eyes absent homolog 1 | *Homo sapiens* | Vertebrata | protein from NCBI | NP_001275503.1 |
| Hv_Eya | eyes absent homolog 1 | *Hydra vulgaris* | Cnidaria | protein from NCBI | XP_012556911.1 |
| La_Eya | eyes absent homolog 1 | *Lingula anatina* | Brachiopoda | protein from NCBI | XP_013418681.1 |
| My_Eya | eyes absent homolog 1 | *Mizuhopecten yessoensis* | Mollusca | protein from NCBI | XP_021379793.1 |
| Mm_Eya1 | eyes absent homolog 1 | *Mus musculus* | Vertebrata | protein from NCBI | NP_001297388.1 |
| Ot_Eya | eyes absent homolog 1 | *Onthophagus taurus* | Arthropoda | protein from NCBI | XP_022912963.1 |
| Pt_Eya | eyes absent homolog 1 | *Parasteatoda tepidariorum* | Arthropoda | protein from NCBI | XP_015924744.1 |
| Pca_Eya | eyes absent homolog 1 | *Pomacea canaliculata* | Mollusca | protein from NCBI | XP_025111355.1 |
| Sp_Eya | eyes absent homolog 1 | *Strongylocentrotus purpuratus* | Echinodermata | protein from NCBI | XP_011666140.1 |
| Aco_Eya | eya | *Ananas comosus* | plants | protein from NCBI | OAY68533.1 |
| Os_Eya | eya | *Oryza sativa* | plants | protein from NCBI | BAA85161.1 |

**Table S1** continued

| **tree label** | **protein** | **species** | **clade** | **source** | **sequence** |
| --- | --- | --- | --- | --- | --- |
| Zm_eya | eya | *Zea mays* | plants | protein from NCBI | AIU50393.1 |
| Sca_Eya | eyes absent homolog 1 | *Schizocardium californicum* | Hemichordata | Lowe transcriptome | MT900898 |
| Ek_Eya | eyes absent homolog 1 | *Euperipatoides kanangrensis* | Onychophora | Janssen transcripome | MT900891 |
| Ofu_Eya | eyes absent homolog 1 | *Owenia fusiformis* | Annelida | Hejnol transcriptome | MT900912 |
| Ph_Eya | eyes absent homolog 1 | *Phoronopsis harmeri* | Phoronida | Hejnol transcriptome | MT900918 |
| Pc_Eya | eyes absent homolog 1 | *Priapulus caudatus* | Priapulida | Hejnol transcriptome | MT900882 |
| Tt_Eya | eyes absent homolog 1 | *Terebratalia transversa* | Brachiopoda | Hejnol transcriptome | MT900859 |

**Table S2** Sequences used for the assessment of *eya* genes orthology

| **Tree label** | **protein** | **species** | **clade** | **Homology** | **source** | **sequence** |
| --- | --- | --- | --- | --- | --- | --- |
| Ace_Six4 | six1 | *Acanthaster planci* | Echinodermata | six1/2 | protein from NCBI | XP_022081304.1 |
| Ap_Six1 | Six1 | *Acanthaster planci* | Echinodermata | six1/2 | protein from NCBI | XP_022081304.1 |
| Am_Six1 | six1 | *Apis mellifera* | Arthropoda | six1/2 | protein from NCBI | XP_006564255.1 |
| Am_Six6 | six6 | *Apis mellifera* | Arthropoda | Six4/5 | protein from NCBI | XP_001120698.3 |
| Ac_Six1a | Six1a | *Aplysia californica* | Mollusca | six1/2 | protein from NCBI | XP_005110051.2 |
| Ac_Six1b | six1b | *Aplysia californica* | Mollusca | six4/5 | protein from NCBI | XP_005107679.2 |
| Aj_Six4 | six4 | *Apostichopus japonicus* | Echinodermata | Six4/5 | protein from NCBI | PIK55397.1 |
| Bf_Six | six | *Branchiostoma floridae* | Chordata | Six3/6 | protein from NCBI | EEN45170.1 |
| Cg_Six1 | six1 | *Crassostrea gigas* | Mollusca | six1/2 | protein from NCBI | EKC19049.1 |
| Cg_Six6 | six6 | *Crassostrea gigas* | Mollusca | Six3/6 | protein from NCBI | XP_011426302.1 |
| Cg_Six4 | six4 | *Crassostrea gigas* | Mollusca | six4/5 | protein from NCBI | EKC40506.1 |
| Dr_Six1a | six1a | *Danio rerio* | Vertebrata | six1a | protein from NCBI | XP_021336244.1 |
| Dr_Six1b | six1b | *Danio rerio* | Vertebrata | six1b | protein from NCBI | NP_996978.1 |
| Dr_Six2 | six2 | *Danio rerio* | Vertebrata | six2 | protein from NCBI | NP_571858.1 |
| Dr_Six3 | six3 | *Danio rerio* | Vertebrata | six3 | protein from NCBI | BAA31752.1 |
| Dr_Six4.1 | Six4.1 | *Danio rerio* | Vertebrata | six4a | protein from NCBI | BAB18513.1 |
| Dr_Six4.2 | six4.2 | *Danio rerio* | Vertebrata | six4b | protein from NCBI | BAB18511.1 |
| Dr_Six5 | six5 | *Danio rerio* | Vertebrata | six5b | protein from NCBI | NP_571795.1 |
| Dr_Six6 | Six6a | *Danio rerio* | Vertebrata | six6 | protein from NCBI | NP_957399.1 |
| Da_Six1 | six1 | *Drosophila arizonae* | Arthropoda | six4/5 | protein from NCBI | XP_017872881.1 |
| Dm_Six | sineoculis | *Drosophila melanogaster* | Arthropoda | six1/2 | protein from NCBI | AAF59260.1 |
| Hs_Six1 | six1 | *Homo sapiens* | Vertebrata | six1 | protein from NCBI | CAA62974.1 |
| Hs_Six2 | six2 | *Homo sapiens* | Vertebrata | six2 | protein from NCBI | AAK16583.1 |
| Hs_Six3 | six3 | *Homo sapiens* | Vertebrata | six3 | protein from NCBI | AAD15753.1 |

**Table S2** continued

| **Tree label** | **protein** | **species** | **clade** | **Homology** | **source** | **sequence** |
| --- | --- | --- | --- | --- | --- | --- |
| Hs_Six4 | six4 | *Homo sapiens* | Vertebrata | six4 | protein from NCBI | BAA86223.1 |
| Hs_Six5 | six5 | *Homo sapiens* | Vertebrata | six5 | protein from NCBI | NP_787071.3 |
| Hs_Six6 | six6 | *Homo sapiens* | Vertebrata | six6 | protein from NCBI | AAH87852.1 |
| Hv_Six6 | Six6 | *Hydra vulgaris* | Cnidaria | six3/6 | protein from NCBI | XP_012566992.1 |
| Hv_Six1 | Six1 | *Hydra vulgaris* | Cnidaria | six4/5 | protein from NCBI | XP_012556487.1 |
| La_Six1a | six1a | *Lingula anatina* | Brachiopoda | six1/2 | protein from NCBI | XP_013414609.1 |
| La_Six6 | six6 | *Lingula anatina* | Brachiopoda | six3/6 | protein from NCBI | XP_013418710.1 |
| La_Six1 | six1 | *Lingula anatina* | Brachiopoda | six4/5 | protein from NCBI | XP_023930449.1 |
| Mr_Six3 | six3 | *Metacrinus rotundus* | Echinodermata | Six3/6 | protein from NCBI | ADE59460.1 |
| My_Six6 | six6 | *Mizuhopecten yessoensis* | Mollusca | Six3/6 | protein from NCBI | OWF43532.1 |
| My_Six1 | six1 | *Mizyhopecten yessoensis* | Mollusca | six1/2 | protein from NCBI | XP_021367620.1 |
| My_Six4 | six4 | *Mizyhopecten yessoensis* | Mollusca | six4/5 | protein from NCBI | OWF54346.1 |
| Mm_Six1 | six1 | *Mus musculus* | Vertebrata | six1 | protein from NCBI | CAA56585.1 |
| Mm_Six2 | six2 | *Mus musculus* | Vertebrata | six2 | protein from NCBI | BAA11825.1 |
| Mm_Six3 | Six3 | *Mus musculus* | Vertebrata | Six3 | protein from NCBI | AAH94426.1 |
| Mm_Six4 | six4 | *Mus musculus* | Vertebrata | six4 | protein from NCBI | NP_035512.1 |
| Mm_Six5 | six5 | *Mus musculus* | Vertebrata | six5 | protein from NCBI | NP_035513.1 |
| Mm_Six6 | six6 | *Mus musculus* | Vertebrata | six6 | protein from NCBI | EDL36517.1 |
| Nv_Six1 | six1/2 | *Nematostella vectensis* | Cnidaria | six1/2 | protein from NCBI | SJX71969.1 |
| Nv_Six3 | six3/6 | *Nematostella vectensis* | Cnidaria | six3/6 | protein from NCBI | AGD98926.1 |
| Nv_Six4 | six4/5 | *Nematostella vectensis* | Cnidaria | six4/5 | protein from NCBI | SJX71970.1 |
| Of_Six1 | six1 | *Orbicella faveolata* | Cnidaria | six1/2 | protein from NCBI | XP_020605799.1 |
| Pt_Six1 | six1 | *Parasteatoda tepidariorum* | Arthropoda | six1/2a | protein from NCBI | XP_015903690.1 |
| Pt_Six1b | six1b | *Parasteatoda tepidariorum* | Arthropoda | six1/2b | protein from NCBI | XP_015927973.1 |
| Pt_Six6 | six6 | *Parasteatoda tepidariorum* | Arthropoda | six3/6 | protein from NCBI | XP_015922109.1 |

**Table S2** continued

| **Tree label** | **protein** | **species** | **clade** | **Homology** | **source** | **sequence** |
| --- | --- | --- | --- | --- | --- | --- |
| Pt_Six3 | six3 | *Parasteatoda tepidariorum* | Arthropoda | six4/5 | protein from NCBI | XP_015927956.1 |
| Pd_Six2 | six2 | *Platynereis dumerilii* | Annelida | six1/2 | protein from NCBI | CAC86663.1 |
| Pd_Six3 | six3 | *Platynereis dumerilii* | Annelida | six3/6 | protein from NCBI | CAR66435.1 |
| Pc_Six4_5 | six1 | *Pomacea canaliculata* | Mollusca | six4/5 | protein from NCBI | XP_025092083.1 |
| Pca_Six6 | six6 | *Priapulus caudatus* | Priapulida | six3/6 | protein from NCBI | XP_014664642.1 |
| Pca_Six1 | six1 | *Priapulus caudatus* | Priapulida | six4/5 | protein from NCBI | XP_014678215.1 |
| Sk_Six1 | six1 | *Saccoglossus kowalevskii* | Hemichordata | six1/2 | protein from NCBI | NP_001277017.1 |
| Sk_Six3 | six3 | *Saccoglossus kowalevskii* | Hemichordata | six3/6 | protein from NCBI | XP_006817024.1 |
| Sc_Six3 | six3 | *Schizocardium californicum* | Hemichordata | Six3/6 | protein from NCBI | ARO85860.1 |
| Spu_Six1 | six1 | *Strongylocentrotus purpuratus* | Echinodermata | six1/2 | protein from NCBI | AEZ53927.1 |
| Spu_Six4 | six4 | *Strongylocentrotus purpuratus* | Echinodermata | Six4/5 | protein from NCBI | XP_781616.2 |
| Tt_Six3_6 | six3/6 | *Terebratalia transversa* | Brachiopoda | Six3/6 | protein from NCBI | AEZ03831.1 |
| Tc_six | sine oculis | *Tribolium castaneum* | Arthropoda | six1/2 | protein from NCBI | EFA04748.2 |
| Tc_optix | optix | *Tribolium castaneum* | Arthropoda | six3/6 | protein from NCBI | NP_001106938.1 |
| Tc_Six1 | six1 | *Tribolium castaneum* | Arthropoda | six4/5 | protein from NCBI | XP_015833917.1 |
| Sca_Six1_2 | six1/2 | *Schizocardium californicum* | Hemichordata | six1/2 | Lowe transcriptome | MT900894 |
| Ek_Six1_2 | six1/2 | *Euperipatoides kanangrensis* | Onychophora | six1/2 | Janssen transcripome | MT900890 |
| Ofu_Six1_2 | six1/2 | *Owenia fusiformis* | Annelida | six1/2 | Hejnol transcriptome | MT900902 |
| Ph_Six1_2 | six1/2 | *Phoronopsis harmeri* | Phoronida | six1/2 | Hejnol transcriptome | MT900917 |
| Pc_Six1_2 | six1/2 | *Priapulus caudatus* | Priapulida | six1/2 | Hejnol transcriptome | MT900881 |
| Tt_Six1_2 | six1/2 | *Terebratalia transversa* | Brachiopoda | six1/2 | Hejnol transcriptome | MT900856 |

**Table S3** Sequences used for the assessment of *POU* genes orthology

| **Tree label** | **protein** | **species** | **clade** | **source** | **sequence** |
| --- | --- | --- | --- | --- | --- |
| Cg_POU2 | POU2 | *Crassostrea gigas* | Mollusca | protein from NCBI | XP_011413738.1 |
| Cg_POU3 | POU3 | *Crassostrea gigas* | Mollusca | protein from NCBI | XP_011441819.1 |
| Cv_POU3 | POU3 | *Crassostrea virginica* | Mollusca | protein from NCBI | XP_022293687.1 |
| Cv_POU4 | POU4 | *Crassostrea virginica* | Mollusca | protein from NCBI | XP_022327775.1 |
| Cv_POU6 | POU6 | *Crassostrea virginica* | Mollusca | protein from NCBI | XP_022308452.1 |
| Dr_POU3F1 | POU3F1 | *Danio rerio* | Vertebrata | protein from NCBI | NP_571236.1 |
| Dr_POU3F2 | POU3F2 | *Danio rerio* | Vertebrata | protein from NCBI | NP_571235.1 |
| Dr_POU3F3B | POU3F3b | *Danio rerio* | Vertebrata | protein from NCBI | NP_571177.2 |
| Dr_POU4F1 | POU4F1 | *Danio rerio* | Vertebrata | protein from NCBI | AAH65954.1 |
| Dr_POU5F1 | POU5F1 | *Danio rerio* | Vertebrata | protein from NCBI | AAH49295.1 |
| Dr_POU6F1 | POU6F1 | *Danio rerio* | Vertebrata | protein from NCBI | NP_571188.1 |
| Hs_POU1F1 | POU1F1 | *Homo sapiens* | Vertebrata | protein from NCBI | EAW68867.1 |
| Hs_POU2F1 | POU2F1 | *Homo sapiens* | Vertebrata | protein from NCBI | AAH03571.1 |
| Hs_POU2F2 | POU2F2 | *Homo sapiens* | Vertebrata | protein from NCBI | AAH06101.1 |
| Hs_POU2F3 | POU2F3 | *Homo sapiens* | Vertebrata | protein from NCBI | NP_055167.2 |
| Hs_POU3F1 | POU3F1 | *Homo sapiens* | Vertebrata | protein from NCBI | NP_002690.3 |
| Hs_POU3F2 | POU3F2 | *Homo sapiens* | Vertebrata | protein from NCBI | NP_005595.2 |
| Hs_POU3F3 | POU3F3 | *Homo sapiens* | Vertebrata | protein from NCBI | NP_006227.1 |
| Hs_POU3F4 | POU3F4 | *Homo sapiens* | Vertebrata | protein from NCBI | NP_000298.3 |
| Hs_POU4F1 | POU4F1 | *Homo sapiens* | Vertebrata | protein from NCBI | NP_006228.3 |
| Hs_POU5F1 | POU5F1 | *Homo sapiens* | Vertebrata | protein from NCBI | AQY77032.1 |
| Hs_POU6F1 | POU6F1 | *Homo sapiens* | Vertebrata | protein from NCBI | XP_024304794.1 |
| La_POU2 | POU2 | *Lingula anatina* | Brachiopoda | protein from NCBI | XP_013414918.2 |
| La_POU4 | POU4 | *Lingula anatina* | Brachiopoda | protein from NCBI | XP_013419009.1 |

**Table S3** continued

| **Tree label** | **protein** | **species** | **clade** | **source** | **sequence** |
| --- | --- | --- | --- | --- | --- |
| La_POU6 | POU6 | *Lingula anatina* | Brachiopoda | protein from NCBI | XP_013388150.1 |
| My_POU2 | POU2 | *Mizuhopecten yessoensis* | Mollusca | protein from NCBI | XP_021370535.1 |
| My_POU4 | POU4 | *Mizuhopecten yessoensis* | Mollusca | protein from NCBI | XP_021340548.1 |
| My_POU6 | POU6 | *Mizuhopecten yessoensis* | Mollusca | protein from NCBI | XP_021354648.1 |
| Mm_PO6F1 | POU1F1 | *Mus musculus* | Vertebrata | protein from NCBI | AAH61213.1 |
| Mm_POU1F1 | POU2F1 | *Mus musculus* | Vertebrata | protein from NCBI | NP_035267.2 |
| Mm_POU2F1 | POU2F2 | *Mus musculus* | Vertebrata | protein from NCBI | AAI05648.1 |
| Mm_POU2F2 | POU2F3 | *Mus musculus* | Vertebrata | protein from NCBI | NP_035269.2 |
| Mm_POU2F3 | POU3F1 | *Mus musculus* | Vertebrata | protein from NCBI | NP_035271.1 |
| Mm_POU3F1 | POU3F2 | *Mus musculus* | Vertebrata | protein from NCBI | NP_032925.1 |
| Mm_POU3F2 | POU3F3 | *Mus musculus* | Vertebrata | protein from NCBI | NP_032926.2 |
| Mm_POU3F3 | POU3F4 | *Mus musculus* | Vertebrata | protein from NCBI | NP_032927.1 |
| Mm_POU3F4 | POU4F3 | *Mus musculus* | Vertebrata | protein from NCBI | NP_620395.2 |
| Mm_POU4F3 | POU5F1 | *Mus musculus* | Vertebrata | protein from NCBI | NP_083591.1 |
| Mm_POU5F1 | POU6F1 | *Mus musculus* | Vertebrata | protein from NCBI | NP_001291894.1 |
| Pt_POU2 | POU2 | *Parasteatoda tepidariorum* | Arthropoda | protein from NCBI | XP_021000349.1 |
| Pt_POU3 | POU3 | *Parasteatoda tepidariorum* | Arthropoda | protein from NCBI | XP_015910265.1 |
| Pt_POU4 | POU4 | *Parasteatoda tepidariorum* | Arthropoda | protein from NCBI | XP_015918995.1 |
| Pt_POU6 | POU6 | *Parasteatoda tepidariorum* | Arthropoda | protein from NCBI | XP_021003103.1 |
| Pc_POU2 | POU2 | *Pomacea canaliculata* | Mollusca | protein from NCBI | XP_025103000.1 |
| Pca_POU4 | POU4 | *Pomacea canaliculata* | Mollusca | protein from NCBI | XP_025087876.1 |
| Pc_POU6 | POU6 | *Pomacea canaliculata* | Mollusca | protein from NCBI | XP_025094150.1 |
| Pca_POU2 | POU2 | *Priapulus caudatus* | Priapulida | protein from NCBI | XP_014675686.1 |
| Pc_POU4 | POU4 | *Priapulus caudatus* | Priapulida | protein from NCBI | XP_014671546.1 |
| Sk_POU3 | POU3 | *Saccoglossus kowalevskii* | Hemichordata | protein from NCBI | NP_001161508.1 |

**Table S3** continued

| **Tree label** | **protein** | **species** | **clade** | **source** | **sequence** |
| --- | --- | --- | --- | --- | --- |
| Tc_POU2 | POU2 | *Tribolium castaneum* | Arthropoda | protein from NCBI | KYB27833.1 |
| Tc_POU6 | POU6 | *Tribolium castaneum* | Arthropoda | protein from NCBI | EFA13130.2 |
| Xc_POU1F1 | POU1F1 | *Xiphophorus couchianus* | Vertebrata | protein from NCBI | XP_027878947.1 |
| Xc_POU2F1 | POU2F1 | *Xiphophorus couchianus* | Vertebrata | protein from NCBI | XP_027879658.1 |
| Xc_POU2F2 | POU2F2 | *Xiphophorus couchianus* | Vertebrata | protein from NCBI | XP_027864127.1 |
| Xc_POU2F3 | POU2F3 | *Xiphophorus couchianus* | Vertebrata | protein from NCBI | XP_027887813.1 |
| Sca_POU3 | POU3 | *Schizocardium californicum* | Hemichordata | Lowe transcriptome | MT900897 |
| Ek_POU3 | POU3 | *Euperipatoides kanangrensis* | Onychophora | Janssen transcripome | MT900892 |
| Ofu_POU3 | POU3 | *Owenia fusiformis* | Annelida | Hejnol transcriptome | MT900911 |
| Ph_POU3 | POU3 | *Phoronopsis harmeri* | Phoronida | Hejnol transcriptome | MT900921 |
| Pc_POU3 | POU3 | *Priapulus caudatus* | Priapulida | Hejnol transcriptome | MT900885 |
| Tt_POU3 | POU3 | *Terebratalia transversa* | Brachiopoda | Hejnol transcriptome | MT900858 |

**Table S4** Sequences used for the assessment of zinc finger genes orthology

| **Tree label** | **protein** | **species** | **clade** | **source** | **sequence** |
| --- | --- | --- | --- | --- | --- |
| Ap_osr1 | OSR1 | *Acanthaster planci* | Echinodermata | protein from NCBI | XP_022102762.1 |
| Ap_osr2 | OSR2 | *Acanthaster planci* | Echinodermata | protein from NCBI | XP_022081172.1 |
| Ap_sall1 | sall 1 | *Acanthaster planci* | Echinodermata | protein from NCBI | XP_022083926.1 |
| Ac_osr2 | OSR2 | *Apis cerana* | Arthropoda | protein from NCBI | XP_016914606.1 |
| Am_hb | hunchback | *Apis mellifera* | Arthropoda | protein from NCBI | AER27698.1 |
| Bb_osr2 | OSR2 | *Branchiostoma belcheri* | Chordata | protein from NCBI | XP_019618804.1 |
| Ce_hb | hunchback | *Caerorhabditis elegans* | Nematoda | protein from NCBI | CCD69456.1 |
| Cc_hb | hunchback | *Capitella capitata* | Annelida | protein from NCBI | AAY43811.1 |
| Cca_sall3 | sall 3 | *Ceratina calcarata* | Arthropoda | protein from NCBI | XP_017889078.1 |
| Cg_hb | hunchback | *Crassostrea gigas* | Mollusca | protein from NCBI | XP_019923756.1 |
| Cg_osr1 | OSR1 | *Crassostrea gigas* | Mollusca | protein from NCBI | XP_011418777.1 |
| Cg_sall1 | sall 1 | *Crassostrea gigas* | Mollusca | protein from NCBI | XP_011449486.1 |
| Cv_osr1 | OSR1 | *Crassostrea virginica* | Mollusca | protein from NCBI | XP_022319722.1 |
| Cv_sall3 | sall 3 | *Crassostrea virginica* | Mollusca | protein from NCBI | XP_022340917.1 |
| Dr_osr1 | OSR1 | *Danio rerio* | Vertebrata | protein from NCBI | NP_001006079.1 |
| Dr_osr2 | OSR2 | *Danio rerio* | Vertebrata | protein from NCBI | NP_001017694.1 |
| Dm_hb | hunchback | *Drosophila melanogaster* | Arthropoda | protein from NCBI | NP_731268.1 |
| Dm_ods | odd skipped | *Drosophila melanogaster* | Arthropoda | protein from NCBI | NP_722922.1 |
| Gb_hb | hunchback | *Gryllus bimaculatus* | Arthropoda | protein from NCBI | BAD12839.1 |
| Hr_hb | hunchback | *Helobdella robusta* | Annelida | protein from NCBI | AAY43810.1 |
| Hs_osr1 | OSR1 | *Homo sapiens* | Vertebrata | protein from NCBI | NP_660303.1 |
| Hs_osr2 | OSR2 | *Homo sapiens* | Vertebrata | protein from NCBI | XP_011515128.1 |
| Hs_sall1 | sall 1 | *Homo sapiens* | Vertebrata | protein from NCBI | EAW82777.1 |
| Hs_sall2 | sall 2 | *Homo sapiens* | Vertebrata | protein from NCBI | NP_001351493.1 |

**Table S4** continued

| **Tree label** | **protein** | **species** | **clade** | **source** | **sequence** |
| --- | --- | --- | --- | --- | --- |
| Hs_sall4 | sall 4 | *Homo sapiens* | Vertebrata | protein from NCBI | NP_065169.1 |
| La_hb | hunchback | *Lingula anatina* | Brachiopoda | protein from NCBI | XP_013420677.1 |
| La_osr2 | OSR2 | *Lingula anatina* | Brachiopoda | protein from NCBI | XP_013400144.2 |
| La_sall | sall | *Lingula anatina* | Brachiopoda | protein from NCBI | XP_013396129.1 |
| My_hb | hunchback | *Mizuhopecten yessoensis* | Mollusca | protein from NCBI | OWF44667.1 |
| My_osr2 | OSR2 | *Mizuhopecten yessoensis* | Mollusca | protein from NCBI | OWF36140.1 |
| My_sall1 | sall 1 | *Mizuhopecten yessoensis* | Mollusca | protein from NCBI | XP_021379224.1 |
| My_sall3 | sall 3 | *Mizuhopecten yessoensis* | Mollusca | protein from NCBI | OWF37915.1 |
| Mm_sall1 | sall 1 | *Mus musculus* | Vertebrata | protein from NCBI | XP_006531301.1 |
| Mm_sall2 | sall 2 | *Mus musculus* | Vertebrata | protein from NCBI | NP_056587.2 |
| Mm_sall3 | sall 3 | *Mus musculus* | Vertebrata | protein from NCBI | XP_006526522.1 |
| Mm_sall4 | sall 4 | *Mus musculus* | Vertebrata | protein from NCBI | NP_780512.2 |
| Oh_sall2 | sall 2 | *Ophiophagus hannah* | Vertebrata | protein from NCBI | ETE70785.1 |
| Oh_sall3 | sall 3 | *Ophiophagus hannah* | Vertebrata | protein from NCBI | ETE67890.1 |
| Oh_sall4 | sall 4 | *Ophiophagus hannah* | Vertebrata | protein from NCBI | ETE66117.1 |
| Pm_sall1 | sall 1 | *Papilio machaon* | Arthropoda | protein from NCBI | KPJ13820.1 |
| Px_sall3 | sall 3 | *Papilio xuthus* | Arthropoda | protein from NCBI | KPI93779.1 |
| Pt_hb | hunchback | *Parasteatoda tepidariorum* | Arthropoda | protein from NCBI | CAX11339.1 |
| Pt_osr2 | OSR2 | *Parasteatoda tepidariorum* | Arthropoda | protein from NCBI | XP_015910072.1 |
| Pca_sall3 | sall 3 | *Pomacea canaliculata* | Mollusca | protein from NCBI | XP_025114875.1 |
| Rn_osr1 | OSR1 | *Rattus norvegicus* | Vertebrata | protein from NCBI | NP_001100186.1 |
| Rn_osr2 | OSR2 | *Rattus norvegicus* | Vertebrata | protein from NCBI | XP_017450351.1 |
| Sk_osr2 | OSR2 | *Saccoglossus kowalevskii* | Hemichordata | protein from NCBI | XP_006814614.1 |
| Sk_sall | Sall | *Saccoglossus kowalevskii* | Hemichordata | protein from NCBI | ALR88687.1-1 |

**Table S4** continued

| **Tree label** | **protein** | **species** | **clade** | **source** | **sequence** |
| --- | --- | --- | --- | --- | --- |
| Sm_hb | hunchback-like | *Schmidtea mediterranea* | Platyhelminthes | protein from NCBI | JN837710.1 |
| Spu_osr2 | OSR2 | *Strongylocentrotus purpuratus* | Echinodermata | protein from NCBI | XP_011678010.1 |
| Sp_sall1 | sall 1 | *Strongylocentrotus purpuratus* | Echinodermata | protein from NCBI | XP_011673616.1 |
| Tc_hb | hunchback | *Tribolium castaneum* | Arthropoda | protein from NCBI | CAA62821.1 |
| Tc_ods | odd skipped | *Tribolium castaneum* | Arthropoda | protein from NCBI | EFA09193.2 |
| Sca_osr | Osr | *Schizocardium californicum* | Hemichordata | Lowe transcriptome | MT900896 |
| Sca_sall | Sall | *Schizocardium californicum* | Hemichordata | Lowe transcriptome | MT900895 |
| Ofu_hb | hunchback | *Owenia fusiformis* | Annelida | Hejnol transcriptome | MT900904 |
| Ofu_osra | OsrA | *Owenia fusiformis* | Annelida | Hejnol transcriptome | MT900909 |
| Ofu_osrb | OsrB | *Owenia fusiformis* | Annelida | Hejnol transcriptome | MT900910 |
| Ofu_sall | Sall | *Owenia fusiformis* | Annelida | Hejnol transcriptome | MT900903 |
| Ph_hb | hunchback | *Phoronopsis harmeri* | Phoronida | Hejnol transcriptome | MT900922 |
| Ph_osr | Osr | *Phoronopsis harmeri* | Phoronida | Hejnol transcriptome | MT900920 |
| Ph_sall | Sall | *Phoronopsis harmeri* | Phoronida | Hejnol transcriptome | MT900919 |
| Pc_hb | hunchback | *Priapulus caudatus* | Priapulida | Hejnol transcriptome | MT900886 |
| Pc_osr | Osr | *Priapulus caudatus* | Priapulida | Hejnol transcriptome | MT900884 |
| Pc_sall | Sall | *Priapulus caudatus* | Priapulida | Hejnol transcriptome | MT900883 |
| Tt_hb | hunchback | *Terebratalia transversa* | Brachiopoda | Hejnol transcriptome | MT900857 |
| Tt_osr | Osr | *Terebratalia transversa* | Brachiopoda | Hejnol transcriptome | MT900861 |
| Tt_sall | Sall | *Terebratalia transversa* | Brachiopoda | Hejnol transcriptome | MT900860 |

**Table S5** Sequences used for the assessment of *lhx* genes orthology

| **Tree label** | **protein** | **species** | **clade** | **source** | **sequence** |
| --- | --- | --- | --- | --- | --- |
| Ap_Lhx5 | Lhx5 | *Acanthaster planci* | Echinodermata | protein from NCBI | XP_022101121.1 |
| Aj_Lhx4 | Lhx4 | *Apostichopus japonicus* | Echinodermata | protein from NCBI | PIK45227.1 |
| Bf_Lhx3 | Lhx3 | *Branchiostoma floridae* | Chordata | protein from NCBI | EEN47850.1 |
| Cc_Lhx3 | Lhx3 | *Ceratitis capitata* | Arthropoda | protein from NCBI | XP_012155171.1 |
| Cg_Lhx3 | Lhx3 | *Crassostrea gigas* | Mollusca | protein from NCBI | XP_011456185.1 |
| Cg_Lhx5 | Lhx5 | *Crassostrea gigas* | Mollusca | protein from NCBI | XP_011450364.1 |
| Cv_Lhx1 | Lhx1 | *Crassostrea virginica* | Mollusca | protein from NCBI | XP_022343409.1 |
| Cv_Lhx9 | Lhx9 | *Crassostrea virginica* | Mollusca | protein from NCBI | XP_022322359.1 |
| De_Lhx1 | lhx1 | *Drosophila erecta* | Arthropoda | protein from NCBI | XP_001977402.1 |
| De_Lhx3 | Lhx3 | *Drosophila erecta* | Arthropoda | protein from NCBI | XP_026836663.1 |
| Dh_apt | apterous | *Drosophila hydei* | Arthropoda | protein from NCBI | XP_023173008.1 |
| Dm_zyxin | zyxin | *Drosophila melanogaster* | Arthropoda | protein from NCBI | NP_652015.1 |
| Dp_Lhx4 | Lhx4 | *Drosophila persimilis* | Arthropoda | protein from NCBI | XP_026843596.1 |
| Ep_Lhx1 | Lhx1 | *Exaiptasia pallida* | Cnidaria | protein from NCBI | XP_020905808.1 |
| Fa_Lhx1 | Lhx1 | *Fopius arisanus* | Arthropoda | protein from NCBI | XP_011308710.1 |
| Fa_Lhx5 | Lhx5 | *Fopius arisanus* | Arthropoda | protein from NCBI | XP_011308720.1 |
| Fa_Lhx9 | Lhx9 | *Fopius arisanus* | Arthropoda | protein from NCBI | JAG84074.1 |
| Hs_Lhx9 | Lhx9 | *Harpegnathos saltator* | Arthropoda | protein from NCBI | EFN77731.1 |
| Hg_Lhx1-5 | Lhx1/5 | *Holothuria glaberrima* | Echinodermata | protein from NCBI | ALI93842.1 |
| Hs_Lhx1 | Lhx1 | *Homo sapiens* | Vertebrata | protein from NCBI | NP_005559.2 |
| Hs_Lhx2 | Lhx2 | *Homo sapiens* | Vertebrata | protein from NCBI | NP_004780.3 |
| Hs_Lhx3 | Lhx3 | *Homo sapiens* | Vertebrata | protein from NCBI | NP_835258.1 |
| Hs_Lhx4 | Lhx4 | *Homo sapiens* | Vertebrata | protein from NCBI | NP_203129.1 |
| Hs_Lhx5 | Lhx5 | *Homo sapiens* | Vertebrata | protein from NCBI | NP_071758.1 |

**Table S5** continued

| **Tree label** | **protein** | **species** | **clade** | **source** | **sequence** |
| --- | --- | --- | --- | --- | --- |
| Hs_Lhx6 | Lhx6 | *Homo sapiens* | Vertebrata | protein from NCBI | NP_001335119.1 |
| Hs_Lhx8 | Lhx8 | *Homo sapiens* | Vertebrata | protein from NCBI | NP_001001933.1 |
| La_Lhx1 | Lhx1 | *Lingula anatina* | Brachiopoda | protein from NCBI | XP_013413189.1 |
| La_Lhx3 | Lhx3 | *Lingula anatina* | Brachiopoda | protein from NCBI | XP_013382228.1 |
| La_Lhx9 | Lhx9 | *Lingula anatina* | Brachiopoda | protein from NCBI | XP_013402764.1 |
| Lg_Lhx1_5 | Lhx1/5 | *Lottia gigantea* | Mollusca | protein from NCBI | BAQ19229.1 |
| My_Lhx1 | Lhx1 | *Mizuhopecten yessoensis* | Mollusca | protein from NCBI | XP_021372779.1 |
| My_Lhx3 | Lhx3 | *Mizuhopecten yessoensis* | Mollusca | protein from NCBI | XP_021340785.1 |
| My_Lhx9 | Lhx9 | *Mizuhopecten yessoensis* | Mollusca | protein from NCBI | OWF35831.1 |
| Mm_zyxin | zyxin | *Mus musculus* | Vertebrata | protein from NCBI | CAA67510.1 |
| Na_Lhx1 | Lhx1 | *Neanthes arenaceodentata* | Annelida | protein from NCBI | AEN75258.1 |
| Nv_Lhx1 | Lhx1 | *Nematostella vectensis* | Cnidaria | protein from NCBI | XP_001631723.2 |
| Pd_Lhx1_5 | Lhx1/5 | *Platynereis dumerilii* | Annelida | protein from NCBI | ADG26732.1 |
| Pc_Lhx3 | Lhx3 | *Pomacea canaliculata* | Mollusca | protein from NCBI | XP_025113261.1 |
| Pc_Lhx9 | Lhx9 | *Pomacea canaliculata* | Mollusca | protein from NCBI | XP_025113346.1 |
| Rn_Lhx1 | Lhx1 | *Rattus norvegicus* | Vertebrata | protein from NCBI | NP_665887.3 |
| Rn_Lhx2 | Lhx2 | *Rattus norvegicus* | Vertebrata | protein from NCBI | NP_001100041.1 |
| Rn_Lhx3 | Lhx3 | *Rattus norvegicus* | Vertebrata | protein from NCBI | XP_001059910.2 |
| Rn_Lhx4 | Lhx4 | *Rattus norvegicus* | Vertebrata | protein from NCBI | NP_001101818.1 |
| Rn_Lhx5 | Lhx5 | *Rattus norvegicus* | Vertebrata | protein from NCBI | NP_620605.1 |
| Rn_Lhx6 | Lhx6 | *Rattus norvegicus* | Vertebrata | protein from NCBI | NP_001101307.1 |
| Rn_Lhx8 | Lhx8 | *Rattus norvegicus* | Vertebrata | protein from NCBI | NP_001012219.1 |
| Rn_Lhx9 | Lhx9 | *Rattus norvegicus* | Vertebrata | protein from NCBI | NP_852032.1 |
| Sk_Lhx1_5 | Lhx1/5 | *Saccoglossus kowalevskii* | Hemichordata | protein from NCBI | XP_002736028.1 |

**Table S5** continued

| **Tree label** | **protein** | **species** | **clade** | **source** | **sequence** |
| --- | --- | --- | --- | --- | --- |
| SPu_Lhx1 | Lhx1 | *Strongylocentrotus purpuratus* | Echinodermata | protein from NCBI | NP_999810.1 |
| Tc_apt | apterous | *Tribolium castaneum* | Arthropoda | protein from NCBI | NP_001139388.1 |
| Tc_Lhx3 | Lhx3 | *Tribolium castaneum* | Arthropoda | protein from NCBI | XP_008193584.2 |
| Tc_Lhx5 | Lhx5 | *Tribolium castaneum* | Arthropoda | protein from NCBI | XP_015836533.1 |
| Xm_Lhx1 | Lhx1 | *Xiphophorus maculatus* | Vertebrata | protein from NCBI | XP_005813915.1 |
| Xm_Lhx2 | Lhx2 | *Xiphophorus maculatus* | Vertebrata | protein from NCBI | XP_023197525.1 |
| Xm_Lhx3 | Lhx3 | *Xiphophorus maculatus* | Vertebrata | protein from NCBI | XP_023193771.1 |
| Xm_Lhx4 | Lhx4 | *Xiphophorus maculatus* | Vertebrata | protein from NCBI | XP_005800587.1 |
| Xm_Lhx5 | Lhx5 | *Xiphophorus maculatus* | Vertebrata | protein from NCBI | XP_023193714.1 |
| Xm_Lhx6 | Lhx6 | *Xiphophorus maculatus* | Vertebrata | protein from NCBI | XP_023193618.1 |
| Xm_Lhx8 | Lhx8 | *Xiphophorus maculatus* | Vertebrata | protein from NCBI | XP_023194965.1 |
| Xm_Lhx9 | Lhx9 | *Xiphophorus maculatus* | Vertebrata | protein from NCBI | XP_023195825.1 |
| Xm_Zyxin | zyxin | *Xiphophorus maculatus* | Vertebrata | protein from NCBI | XP_005798910.2 |
| Sca_Lhx1_5 | Lhx1/6 | *Schizocardium californicum* | Hemichordata | protein from NCBI | ARO85855.1 |
| Gala_Lhx1_a | Lhx1 | *Galathowenia sp.* | Annelida | Hejnol transcriptome | MT900874 |
| Gala_Lhx1_b | Lhx1 | *Galathowenia sp.* | Annelida | Hejnol transcriptome | MT900875 |
| Gala_Lhx1_c | Lhx1 | *Galathowenia sp.* | Annelida | Hejnol transcriptome | MT900876 |
| Ofu_Lhx1_a | Lhx1/5a | *Owenia fusiformis* | Annelida | Hejnol transcriptome | MT900905 |
| Ofu_Lhx1_b | Lhx1/5b | *Owenia fusiformis* | Annelida | Hejnol transcriptome | MT900906 |
| Ofu_Lhx1_c | Lhx1/5c | *Owenia fusiformis* | Annelida | Hejnol transcriptome | MT900907 |
| Ofu_Lhx1_d | Lhx1/5d | *Owenia fusiformis* | Annelida | Hejnol transcriptome | MT900908 |
| Ph_Lhx1 | Lhx1/5 | *Phoronopsis harmeri* | Phoronida | Hejnol transcriptome | MT900916 |
| Pc_Lhx1_5 | Lhx1/5 | *Priapulus caudatus* | Priapulida | Hejnol transcriptome | MT900880 |
| Tt_Lhx1_5 | Lhx1/5 | *Terebratalia transversa* | Brachiopoda | Hejnol transcriptome | MT900865 |

**Table S6** Sequences used for the assessment of *ZO1* genes orthology

| **Tree label** | **protein** | **species** | **clade** | **source** | **sequence** |
| --- | --- | --- | --- | --- | --- |
| Hs_ZO1 | ZO1 | *Homo sapiens* | Vertebrata | NP_003248.3 | protein from NCBI |
| Hs_ZO2 | ZO2 | *Homo sapiens* | Vertebrata | AAH27592.1 | protein from NCBI |
| Hs_ZO3 | ZO3 | *Homo sapiens* | Vertebrata | NP_001254490.1 | protein from NCBI |
| Ap_ZO3 | ZO3 | *Acanthaster planci* | Echinodermata | XP_022099789.1 | protein from NCBI |
| Ap_ZO1 | ZO1 | *Acanthaster planci* | Echinodermata | XP_022099788.1 | protein from NCBI |
| Cv_ZO1 | ZO1 | *Crassostrea virginica* | Mollusca | XP_022306903.1 | protein from NCBI |
| Dr_ZO2 | ZO2 | *Danio rerio* | Vertebrata | NP_001188500.1 | protein from NCBI |
| Dr_ZO3 | ZO3 | *Danio rerio* | Vertebrata | NP_001315269.1 | protein from NCBI |
| Dr_ZO1 | ZO1 | *Danio rerio* | Vertebrata | XP_009296416.1 | protein from NCBI |
| Hv_ZO1 | ZO1 | *Hydra vulgaris* | Cnidaria | AAK28322.1 | protein from NCBI |
| Sp_ZO1 | ZO1 | *Stylophora pistillata* | Cnidaria | AAK28322.1 | protein from NCBI |
| Hv_ZO2 | ZO2 | *Hydra vulgaris* | Cnidaria | NP_001296679.1 | protein from NCBI |
| My_ZO1 | ZO1 | *Mizuhopecten yessoensis* | Mollusca | OWF44151.1 | protein from NCBI |
| Tc_ZO2 | ZO2 | *Tribolium castaneum* | Arthropoda | XP_015837878.1 | protein from NCBI |
| Tc_ZO1 | ZO1 | *Tribolium castaneum* | Arthropoda | XP_015837877.1 | protein from NCBI |
| Mm_ZO1 | ZO1 | *Mus musculus* | Vertebrata | NP_033412.2 | protein from NCBI |
| Mm_ZO2 | ZO2 | *Mus musculus* | Vertebrata | AAD19964.1 | protein from NCBI |
| Mm_ZO3 | ZO3 | *Mus musculus* | Vertebrata | NP_001269025.1 | protein from NCBI |
| La_ZO1 | ZO1 | *Lingula anatina* | Brachiopoda | XP_013406126.2 | protein from NCBI |
| Hs_DLG1 | DLG1 | *Homo sapiens* | Vertebrata | ABQ66269.1 | protein from NCBI |
| Hv_DLG1 | DLG1 | *Hydra vulgaris* | Cnidaria | CDG70252.1 | protein from NCBI |
| Dm_DLG1 | DLG1 | *Drospohila melanogater* | Arthropoda | AAQ01226.1 | protein from NCBI |
| Dh_ZO1 | ZO1 | *Drosophila hydei* | Arthropoda | XP_023173757.1 | protein from NCBI |
| Fa_ZO1 | ZO1 | *Fopius arisanus* | Arthropoda | XP_011296991.1 | protein from NCBI |

**Table S6** continued

| **Tree label** | **protein** | **species** | **clade** | **source** | **sequence** |
| --- | --- | --- | --- | --- | --- |
| Bm_ZO1 | ZO1 | *Bombyx mori* | Arthropoda | XP_012550292.1 | protein from NCBI |
| Sp_ZO1 | ZO1 | *Strongylocentrotus purpuratus* | Echinodermata | XP_011663807.1 | protein from NCBI |
| Aj_ZO1 | ZO1 | *Apostichopus japonicus* | Echinodermata | PIK39471.1 | protein from NCBI |
| Sk_ZO1 | ZO1 | *Saccoglossus kowalevskii* | Hemichordata | XP_006813237.1 | protein from NCBI |
| Sca_ZO1 | ZO1 | *Schizocardium californicum* | Hemichordata | Lowe transcriptome | MT900925 |
| Pc_ZO1 | ZO1 | *Priapulus caudatus* | Priapulida | Hejnol transcriptome | MT900879 |
| Ph_ZO1 | ZO1 | *Phoronopsis harmeri* | Phoronida | Hejnol transcriptome | MT900915 |
| Ofu_ZO1 | ZO1 | *Owenia fusiformis* | Annelida | Hejnol transcriptome | MT900901 |
| Tt_ZO1 | ZO1 | *Terebratalia transversa* | Brachiopoda | Hejnol transcriptome | MT900864 |
| Ek_ZO1 | ZO1 | *Euperipatoides kanangrensis* | Onychophora | Janssen transcripome | MT900889 |

**Table S7** Sequences used for the assessment of *nephrin* and *kirre* genes orthology

| **Tree label** | **protein** | **species** | **clade** | **source** | **sequence** | **transcriptome** |
| --- | --- | --- | --- | --- | --- | --- |
| Bbe_NPHNA | Branchiostoma belcheri nephrin a | *Branchiostoma belcheri* | Chordata | public transcriptome | XM_019771836.1 | PRJNA358734 |
| Bbe_NPHNB | Branchiostoma belcheri nephrin b | *Branchiostoma belcheri* | Chordata | public transcriptome | XM_019792173.1 | PRJNA358734 |
| Bbe_kirre | Branchiostoma belcheri kirre | *Branchiostoma belcheri* | Chordata | public transcriptome | XM_019771840.1 | PRJNA358734 |
| Bfl_kirre | Branchiostoma floridae kirre | *Branchiostoma floridae* | Chordata | public transcriptome | GETA01024979.1 | PRJNA215261 |
| Bfl_NPHN | Branchiostoma floridae nephrin | *Branchiostoma floridae* | Chordata | public transcriptome | GETA01026506.1 | PRJNA215261 |
| Cnem_NPHN1 | Cepaea nemoralis nephrin a | *Cepaea nemoralis* | Mollusca | public transcriptome | GFLU01131598.1 | PRJNA377398 |
| Cnem_NPHN2 | Cepaea nemoralis nephrin b | *Cepaea nemoralis* | Mollusca | public transcriptome | GFLU01067355.1 | PRJNA377398 |
| Cnem_kirre | Cepaea nemoralis kirre | *Cepaea nemoralis* | Mollusca | public transcriptome | GFLU01084564.1 | PRJNA377398 |
| Cin_NPHN | Ciona intestinalis nephrin | *Ciona intestinalis* | Chordata | public transcriptome | XM_002122711.5 | PRJNA187185 |
| Cin_kirre | Ciona intestinalis kirre | *Ciona intestinalis* | Chordata | public transcriptome | XM_018813641.2 | PRJNA187185 |
| Csav_NPHN | Ciona savignyi nephrin | *Ciona savignyi* | Chordata | public transcriptome | GGEI01054665.1 | PRJNA427851 |
| Csav_kirre | Ciona savignyi kirre | *Ciona savignyi* | Chordata | public transcriptome | GGEI01079964.1 | PRJNA427851 |
| Dre_kirreA | Danio rerio kirre a | *Danio rerio* | Chordata | public transcriptome | GDQH01015268.1 | PRJNA280983 |
| Dre_kirreB | Danio rerio kirre b | *Danio rerio* | Chordata | public transcriptome | GDQH01000295.1 | PRJNA280983 |
| Dre_kirreC | Danio rerio kirre c | *Danio rerio* | Chordata | public transcriptome | GDQH01020935.1 | PRJNA280983 |
| Dmag_NPHN | Daphnia magna nephrin | *Daphnia magna* | Arthropoda | public transcriptome | GDIQ01059934.1 | PRJNA284518 |
| Dmag_kirre | Daphnia magna kirre | *Daphnia magna* | Arthropoda | public transcriptome | GDIP01030495.1 | PRJNA284518 |
| Gpau_NPHN | Glossoscolex paulistus nephrin | *Glossoscolex paulistus* | Annelida | public transcriptome | GBIL01045480.1 | PRJNA253195 |
| Gpau_kirre | Glossoscolex paulistus kirre | *Glossoscolex paulistus* | Annelida | public transcriptome | GBIL01044598.1 | PRJNA253195 |
| Gbim_NPHN | Gryllus bimaculatus nephrin | *Gryllus bimaculatus* | Arthropoda | public transcriptome | GFMG01297029.1 | PRJNA376023 |
| Gbim_kirre | Gryllus bimaculatus kirre | *Gryllus bimaculatus* | Arthropoda | public transcriptome | GFMG01177842.1 | PRJNA376023 |
| Gpr_NPHN | Gymnocypris przewalskii nephrin | *Gymnocypris przewalskii* | Chordata | public transcriptome | GEGT01008240.1 | PRJNA257537 |
| Gpr_kirre | Gymnocypris przewalskii kirre | *Gymnocypris przewalskii* | Chordata | public transcriptome | GEGT01017857.1 | PRJNA257537 |
| Hma_NPHN | Hapalochlaena maculosa nephrin | *Hapalochlaena maculosa* | Mollusca | public transcriptome | GEXH01077752.1 | PRJNA337893 |

**Table S7** continued

| **Tree label** | **protein** | **species** | **clade** | **source** | **sequence** | **transcriptome** |
| --- | --- | --- | --- | --- | --- | --- |
| Hma_kirre | Hapalochlaena maculosa kirre | *Hapalochlaena maculosa* | Mollusca | public transcriptome | GEXH01060848.1 | PRJNA337893 |
| Hduj_NPHN | Hypsibius dujardini nephrin | *Hypsibius dujardini* | Tardigrada | public transcriptome | GFGW01008015.1 | PRJNA369152 |
| Hduj_kirre | Hypsibius dujardini kirre | *Hypsibius dujardini* | Tardigrada | public transcriptome | GFGW01003856.1 | PRJNA369152 |
| Lsat_NPHN | Lamellibranchia satsuma nephrin | *Lamellibranchia satsuma* | Annelida | public transcriptome | GEHO01085856.1 | PRJNA231974 |
| Lsat__kirre | Lamellibranchia satsuma kirre | *Lamellibranchia satsuma* | Annelida | public transcriptome | GEHO01059961.1 | PRJNA231974 |
| Pte_kirre | Parasteatoda tepidariorum kirre | *Parasteatoda tepidariorum* | Arthropoda | public transcriptome | IACA01109688.1 | PRJDB4545 |
| Pte_NPHNA | Parasteatoda tepidariorum nephrin a | *Parasteatoda tepidariorum* | Arthropoda | public transcriptome | IACA01109117.1 | PRJDB4545 |
| Pte_NPHNB | Parasteatoda tepidariorum nephrin b | *Parasteatoda tepidariorum* | Arthropoda | public transcriptome | IACA01104794.1 | PRJDB4545 |
| Rma_kirreA | Rhinella marina kirre a | *Rhinella marina* | Chordata | public transcriptome | GFMT01052080.1 | PRJNA383966 |
| Rma_kirreB | Rhinella marina kirre b | *Rhinella marina* | Chordata | public transcriptome | GFMT01061815.1 | PRJNA383966 |
| Rma_kirreC | Rhinella marina kirre c | *Rhinella marina* | Chordata | public transcriptome | GFMT01025395.1 | PRJNA383966 |
| Rma_NPHN | Rhinella marina neprhin | *Rhinella marina* | Chordata | public transcriptome | GFMT01039448.1 | PRJNA383966 |
| Rsor_NPHN | Rotaria sordida nephrin | *Rotaria sordida* | Rotifera | public transcriptome | GDRH01028786.1 | PRJNA295488 |
| Rren_kirre | Rotylenchulus reniformis kirre | *Rotylenchulus reniformis* | Nematoda | public transcriptome | GGVV01029874.1 | PRJNA286314 |
| Rren_NPHNA | Rotylenchulus reniformis nephrin a | *Rotylenchulus reniformis* | Nematoda | public transcriptome | GGVO01056561.1 | PRJNA286314 |
| Rren_NPHNB | Rotylenchulus reniformis nephrin b | *Rotylenchulus reniformis* | Nematoda | public transcriptome | GGVQ01129585.1 | PRJNA286314 |
| Ssa_kirreA | Salmo salar kirre a | *Salmo salar* | Chordata | public transcriptome | GBRB01037564.1 | PRJNA260929 |
| Ssa_kirreB | Salmo salar kirre b | *Salmo salar* | Chordata | public transcriptome | GEGX01274722.1 | PRJNA260929 |
| Ssa_kirreC | Salmo salar kirre c | *Salmo salar* | Chordata | public transcriptome | GGAQ01014876.1 | PRJNA419712 |
| Ssa_kirreD | Salmo salar kirre d | *Salmo salar* | Chordata | public transcriptome | GGAQ01036027.1 | PRJNA419712 |
| Ssa_kirreE | Salmo salar kirre e | *Salmo salar* | Chordata | public transcriptome | GEGX01274715.1 | PRJNA260929 |
| Ssa_kirreF | Salmo salar kirre f | *Salmo salar* | Chordata | public transcriptome | GGAQ01019683.1 | PRJNA419712 |
| Ssa_NPHN | Salmo salar nephrin | *Salmo salar* | Chordata | public transcriptome | GBRB01033467.1 | PRJNA260929 |
| Sth_kirre | Salpa thompsoni kirre | *Salpa thompsoni* | Chordata | public transcriptome | GFCC01107525.1 | PRJNA279245 |

**Table S7** continued

| **Tree label** | **protein** | **species** | **clade** | **source** | **sequence** | **transcriptome** |
| --- | --- | --- | --- | --- | --- | --- |
| Sth_NPHN | Salpa thompsoni nephrin | *Salpa thompsoni* | Chordata | public transcriptome | GFCC01118334.1 | PRJNA279245 |
| Sph_kirre | Sepia pharaonis kirre | *Sepia pharaonis* | Mollusca | public transcriptome | GEIE01010279.1 | PRJNA305947 |
| Sph_NPHN | Sepia pharaonis nephrin | *Sepia pharaonis* | Mollusca | public transcriptome | GEIE01013149.1 | PRJNA305947 |
| Slam_NPHN | Spirobranchus lamarcki nephrin | *Spirobranchus lamarcki* | Annelida | public transcriptome | GGGS01226366.1 | PRJNA433343 |
| Tse_kirre | Tityus serrulatus kirre | *Tityus serrulatus* | Arthropoda | public transcriptome | GBZU01009324.1 | PRJNA261053 |
| Tse_NPHN | Tityus serrulatus nephrin | *Tityus serrulatus* | Arthropoda | public transcriptome | GBZU01019874.1 | PRJNA261053 |
| Tcas_kirre | Tribolium castaneum kirre | *Tribolium castaneum* | Arthropoda | public transcriptome | XM_008192779.2 | PRJNA15718 |
| Tcas_NPHN | Tribolium castaneum nephrin | *Tribolium castaneum* | Arthropoda | public transcriptome | XM_015977663.1 | PRJNA15718 |
| Vin_kirre | Vampirotheutis infernalis kirre | *Vampirotheutis infernalis* | Mollusca | public transcriptome | GGNA01052201.1 | PRJNA342927 |
| Vin_NPHN | Vampirotheutis infernalis nephrin | *Vampirotheutis infernalis* | Mollusca | public transcriptome | GGNA01064433.1 | PRJNA342927 |
| Aca_NPHN | Aplysia califronica nephrin | *Aplysia califronica* | Mollusca | protein from NCBI | XP_012942646.1 | n/a |
| Aca_kirre | Aplysia califronica kirre | *Aplysia califronica* | Mollusca | protein from NCBI | XP_005094996.2 | n/a |
| Bpli_NPHN | Brachionus plicatilis nephrin | *Brachionus plicatilis* | Rotifera | protein from NCBI | RNA22476.1 | n/a |
| Cel_Syg-2 | Caenorhabditis elegans Syg-2 | *Caenorhabditis elegans* | Nematoda | protein from NCBI | NP_001309467.1 | n/a |
| Cel_Syg-1 | Caenorhabditis elegans Syg-1 | *Caenorhabditis elegans* | Nematoda | protein from NCBI | CCD72283.1 | n/a |
| Crem_Syg-2 | Caenorhabditis remanei Syg-2 | *Caenorhabditis remanei* | Nematoda | protein from NCBI | XP_003106204.1 | n/a |
| Crem__Syg-1 | Caenorhabditis remanei Syg-1 | *Caenorhabditis remanei* | Nematoda | protein from NCBI | XP_003109504.1 | n/a |
| Csc_NPHNA | Centruroides sculpturatus nephrin a | *Centruroides sculpturatus* | Arthropoda | protein from NCBI | XP_023238277.1 | n/a |
| Csc_NPHNB | Centruroides sculpturatus nephrin b | *Centruroides sculpturatus* | Arthropoda | protein from NCBI | XP_023231487.1 | n/a |
| Csc_kirre | Centruroides sculpturatus kirre | *Centruroides sculpturatus* | Arthropoda | protein from NCBI | XP_023220646.1 | n/a |
| Clec_NPHN | Cimex lectularis nephrin | *Cimex lectularis* | Arthropoda | protein from NCBI | XP_024084429.1 | n/a |
| Clec_kirre | Cimex lectularis kirre | *Cimex lectularis* | Arthropoda | protein from NCBI | XP_014251757.2 | n/a |
| Cgi_NPHN | Crassostrea gigas nephrin | *Crassostrea gigas* | Mollusca | protein from NCBI | XP_011422664.1 | n/a |
| Cgi_kirre | Crassostrea gigas kirre | *Crassostrea gigas* | Mollusca | protein from NCBI | XP_011422681.1 | n/a |
| Dre_NPHN | Danio rerio nephrin | *Danio rerio* | Chordata | protein from NCBI | NP_001035777.1 | n/a |

**Table S7** continued

| **Tree label** | **protein** | **species** | **clade** | **source** | **sequence** | **transcriptome** |
| --- | --- | --- | --- | --- | --- | --- |
| Dmel_hibris | Drosophila melanogaster hibris | *Drosophila melanogaster* | Arthropoda | protein from NCBI | AAF19446.1 | n/a |
| Dmel_SNS | Drosophila melanogaster SNS | *Drosophila melanogaster* | Arthropoda | protein from NCBI | AAF77184.1 | n/a |
| Dmel_rst | Drosophila melanogaster roughest | *Drosophila melanogaster* | Arthropoda | protein from NCBI | AAF45845.2 | n/a |
| Dmel_kirre | Drosophila melanogaster kirre | *Drosophila melanogaster* | Arthropoda | protein from NCBI | AAF86308.1 | n/a |
| Gga_kirre1 | Gallus gallus kin of irre like 1 | *Gallus gallus* | Chordata | protein from NCBI | XP_423078.3 | n/a |
| Gga_kirre_3 | Gallus gallus kin of irre like 3 | *Gallus gallus* | Chordata | protein from NCBI | XP_004948016.1 | n/a |
| Hsa_NPHN | Homo sapiens nephrin | *Homo sapiens* | Chordata | protein from NCBI | AAG17141.1 | n/a |
| Hsa_kirre1 | Homo sapiens kin of irre like 1 | *Homo sapiens* | Chordata | protein from NCBI | NP_060710.3 | n/a |
| Hsa_kirre2 | Homo sapiens kin of irre like 2 | *Homo sapiens* | Chordata | protein from NCBI | NP_115499.5 | n/a |
| Hsa_kirre3 | Homo sapiens kin of irre like 3 | *Homo sapiens* | Chordata | protein from NCBI | NP_115920.1 | n/a |
| Lch_NPHN | Latimeria chalumnae nephrin | *Latimeria chalumnae* | Chordata | protein from NCBI | XP_014345801.1 | n/a |
| Lch_kirre_1 | Latimeria chalumnae kirre 1 | *Latimeria chalumnae* | Chordata | protein from NCBI | XP_005993211.1 | n/a |
| Lch_kirre_2 | Latimeria chalumnae kirre 2 | *Latimeria chalumnae* | Chordata | protein from NCBI | XP_005999235.1 | n/a |
| Lch_kirre_3 | Latimeria chalumnae kirre 3 | *Latimeria chalumnae* | Chordata | protein from NCBI | XP_014340394.1 | n/a |
| Lana_NPHN | Lingula anatina nephrin | *Lingula anatina* | Brachiopoda | protein from NCBI | XP_013380213.1 | n/a |
| Lana_kirre | Lingula anatina kirre | *Lingula anatina* | Brachiopoda | protein from NCBI | XP_013380223.1 | n/a |
| Mye_NPHN | Mizuhopecten yessoensis nephrin | *Mizuhopecten yessoensis* | Mollusca | protein from NCBI | OWF40059.1 | n/a |
| Mye_kirre | Mizuhopecten yessoensis kirre | *Mizuhopecten yessoensis* | Mollusca | protein from NCBI | OWF40054.1 | n/a |
| Mmu_NPHN | Mus musculus nephrin | *Mus musculus* | Chordata | protein from NCBI | NP_062332.2 | n/a |
| Mmu_kirre1 | Mus musculus kin of irre like 1 | *Mus musculus* | Chordata | protein from NCBI | NP_001164456.1 | n/a |
| Mmu_kirre2 | Mus musculus kin of irre like 2 | *Mus musculus* | Chordata | protein from NCBI | EDL24020.1 | n/a |
| Mmu_kirre3 | Mus musculus kin of irre like 3 | *Mus musculus* | Chordata | protein from NCBI | EDL25368.1 | n/a |
| Obi_NPHN | Octopus bimaculoides nephrin | *Octopus bimaculoides* | Mollusca | protein from NCBI | XP_014776688 | n/a |
| Obi_kirre | Octopus bimaculoides kirre | *Octopus bimaculoides* | Mollusca | protein from NCBI | XP_014783579 | n/a |
| Pdom_kirre | Polistes dominula kirre | *Polistes dominula* | Arthropoda | protein from NCBI | XP_015186955.1 | n/a |

**Table S7** continued

| **Tree label** | **protein** | **species** | **clade** | **source** | **sequence** | **transcriptome** |
| --- | --- | --- | --- | --- | --- | --- |
| Pdom_NPHNA | Polistes dominula nephrin | *Polistes dominula* | Arthropoda | protein from NCBI | XP_015186947.1 | n/a |
| Rvar_kirre | Ramazzottius varieornatus kirre | *Ramazzottius varieornatus* | Tardigrada | protein from NCBI | GAU88376.1 | n/a |
| Rvar_NPHN | Ramazzottius varieornatus nephrin | *Ramazzottius varieornatus* | Tardigrada | protein from NCBI | GAU98753.1 | n/a |
| Sko_kirre | Saccoglossus kovalevskii kirre | *Saccoglossus kovalevskii* | Hemichordata | protein from NCBI | NP_001164704.1 | n/a |
| Sko_NPHN | Saccoglossus kovalevskii nephrin | *Saccoglossus kovalevskii* | Hemichordata | protein from NCBI | XP_006819647.1 | n/a |
| Spu_kirre | Strongylocentrotus purpuratus kirre | *Strongylocentrotus purpuratus* | Echinodermata | protein from NCBI | XP_003723913.1 | n/a |
| Spu_NPHN | Strongylocentrotus purpuratus nephrin | *Strongylocentrotus purpuratus* | Echinodermata | protein from NCBI | XP_011669820.1 | n/a |
| Tnel_kirre | Trichinella nelsoni kirre | *Trichinella nelsoni* | Nematoda | protein from NCBI | KRX18463.1 | n/a |
| Tnel_NPHN | Trichinella nelsoni nephrin | *Trichinella nelsoni* | Nematoda | protein from NCBI | KRX27120.1 | n/a |
| Dere_NCAM1 | NCAM1 | *Drosophila erecta* | Arthropoda | protein from NCBI | XP_001968921.2 | n/a |
| Tcas_NCAM2 | NCAM2 | *Tribolium castaneum* | Arthropoda | protein from NCBI | EFA06824.2 | n/a |
| Hsal_NCAM1 | NCAM1 | *Harpegnathos saltator* | Arthropoda | protein from NCBI | EFN85799.1 | n/a |
| Amel_NCAM2 | NCAM2 | *Apis mellifera* | Arthropoda | protein from NCBI | XP_026296583.1 | n/a |
| Tcas_NCAM1 | NCAM1 | *Tribolium castaneum* | Arthropoda | protein from NCBI | EFA06826.2 | n/a |
| Dmel_Turtle | Turtle | *Drosophila melanogaster* | Arthropoda | protein from NCBI | NP_001303308.1 | n/a |
| Mmu_Turtle | Turtle | *Mus musculus* | Chordata | protein from NCBI | NP_001123259.1 | n/a |
| Hsa_Turtle | Turtle | *Homo sapiens* | Chordata | protein from NCBI | NP_001128522.1 | n/a |
| Hsa_Robo_1 | Robo 1 | *Homo sapiens* | Chordata | protein from NCBI | NP_002932.1 | n/a |
| Dmel_Robo_1 | Robo 1 | *Drosophila melanogaster* | Arthropoda | protein from NCBI | AAM71113.3 | n/a |
| Mmu_Robo_1 | Robo 1 | *Mus musculus* | Chordata | protein from NCBI | NP_062286.2 | n/a |
| Cgi_NCAM1 | NCAM1 | *Crassostrea gigas* | Mollusca | protein from NCBI | EKC41778.1 | n/a |
| Spis_NCAM1 | NCAM1 | *Stylophora pistillata* | Cnidaria | protein from NCBI | PFX24085.1 | n/a |
| Hsa_NCAM1 | NCAM1 | *Homo sapiens* | Chordata | protein from NCBI | NP_000606.3 | n/a |
| Dre_NCAM1 | NCAM1 | *Danio rerio* | Chordata | protein from NCBI | NP_571277.2 | n/a |

**Table S7** continued

| **Tree label** | **protein** | **species** | **clade** | **source** | **sequence** | **transcriptome** |
| --- | --- | --- | --- | --- | --- | --- |
| Scal_kirre | Schizocardium californicum kirre | *Schizocardium californicum* | Hemichordata | Lowe transcriptome | MT900924 | n/a |
| Scal_NPHN | Schizocardium californicum nephrin | *Schizocardium californicum* | Hemichordata | Lowe transcriptome | MT900893 | n/a |
| Ek_NPHN | Euperipatoides kanangrensis nephrin | *Euperipatoides kanangrensis* | Onychophora | Janssen transcripome | MT900887 | n/a |
| Ek_kirre | Euperipatoides kanangrensis kirre | *Euperipatoides kanangrensis* | Onychophora | Janssen transcripome | MT900888 | n/a |
| Hsp_NPHN | Halicryptus spinulosus nephrin | *Halicryptus spinulosus* | Priapulida | Hejnol transcriptome | MT900866 | n/a |
| Hsp_kirre | Halicryptus spinulosus kirre | *Halicryptus spinulosus* | Priapulida | Hejnol transcriptome | MT900923 | n/a |
| Lvi_NPHN | Linneus viridis nephrin | *Linneus viridis* | Nemertea | Hejnol transcriptome | MT900867 | n/a |
| Lvi_kirre | Linneus viridis kirre | *Linneus viridis* | Nemertea | Hejnol transcriptome | MT900868 | n/a |
| Mmem_NPHN | Membranipora membranacea neprhin | *Membranipora membranacea* | Bryozoa | Hejnol transcriptome | MT900873 | n/a |
| Nano_NPHN | Novocrania anomala nephrin | *Novocrania anomala* | Brachiopoda | Hejnol transcriptome | MT900869 | n/a |
| Nano_kirre | Novocrania anomala kirre | *Novocrania anomala* | Brachiopoda | Hejnol transcriptome | MT900870 | n/a |
| Ofu_NPHN | Owenia fusiformis nephrin | *Owenia fusiformis* | Annelida | Hejnol transcriptome | MT900899 | n/a |
| Ofu_kirre | Owenia fusiformis kirre | *Owenia fusiformis* | Annelida | Hejnol transcriptome | MT900900 | n/a |
| Pha_kirre | Phoronopsis harmeri kirre | *Phoronopsis harmeri* | Phoronida | Hejnol transcriptome | MT900914 | n/a |
| Pha_NPHN | Phoronopsis harmeri nephrin | *Phoronopsis harmeri* | Phoronida | Hejnol transcriptome | MT900913 | n/a |
| Pvul_kirre | Pontonema vulgare kirre | *Pontonema vulgare* | Nematoda | Hejnol transcriptome | MT900872 | n/a |
| Pvul_NPHN | Pontonema vulgare nephrin | *Pontonema vulgare* | Nematoda | Hejnol transcriptome | MT900871 | n/a |
| Pc_kirre | Priapulus caudatus kirre | *Priapulus caudatus* | Priapulida | Hejnol transcriptome | MT900878 | n/a |
| Pc_NPHN | Priapulus caudatus nephrin | *Priapulus caudatus* | Priapulida | Hejnol transcriptome | MT900877 | n/a |
| Tt_kirre | Terebratali transversa kirre | *Terebratalia transversa* | Brachiopoda | Hejnol transcriptome | MT900863 | n/a |
| Tt_NPHN | Terebratali transversa nephrin | *Terebratalia transversa* | Brachiopoda | Hejnol transcriptome | MT900862 | n/a |
